# Integrative modeling of guanylate binding protein dimers

**DOI:** 10.1101/2022.12.20.521180

**Authors:** Wibke Schumann, Jennifer Loschwitz, Jens Reiners, Daniel Degrandi, Klaus Pfeffer, Kai Stühler, Gereon Poschmann, Sander H.J. Smits, Birgit Strodel

**Affiliations:** Institute of Theoretical and Computational Chemistry, Heinrich Heine University Düsseldorf, 40225 Düsseldorf, Germany; Institute of Biological Information Processing: Structural Biochemistry, Forschungszentrum Jülich, 52428 Jülich, Germany; Center for Structural Studies Heinrich-Heine-Universität Düsseldorf, 40225 Düsseldorf, Germany; Institute of Medical Microbiology and Hospital Hygiene, Heinrich Heine University, Duesseldorf, Germany; Institute of Molecular Medicine, Proteome Research, Medical Faculty and University Hospital Düsseldorf, Heinrich-Heine University Düsseldorf, Universitätsstraße 1, 40225 Düsseldorf, Germany; Molecular Proteomics Laboratory, Biomedical Research Centre (BMFZ), Heinrich-Heine University Düsseldorf, Universitätsstraße 1, 40225 Düsseldorf, Germany; Institute für Biochemie, Heinrich-Heine-Universität Düsseldorf, 40225 Düsseldorf, Germany

**Author notes:** Contributed equally to this work (Wibke Schumann, Jennifer Loschwitz).

**Keywords:** guanylate binding proteins, protein–protein docking, MD simulation, small angle X-ray scattering, crosslinking mass spectrometry

## Abstract

Guanylate binding proteins (GBPs) are interferon-*γ*-activated large GTPases, effective against intracellular pathogens like *Toxoplasma gondii*. Their host-protective functions require oligomerization, however, the oligomer structures have not been completely resolved yet. Here, we provide dimer models for hGBP1 and the murine GBPs 2 and 7 (mGBP2 and mGBP7) based on integrative modeling that involves the crystal structure of the G domain dimer of hGBP1, cross-linking mass spectrometry (XL-MS), small angle X-ray scattering (SAXS), protein-protein docking, and molecular dynamics (MD) simulations of hGBP1, mGBP2 and mGBP7. We first compare the sequences and protein dynamics of the monomeric hGBP1, mGBP2, and mGBP7, finding that the M/E domain of all three proteins is highly mobile featuring a hinge movement, yet this motion is less pronounced in mGBP7 while its GTPase (G) domain is more flexible. These differences can be explained by the variations in the sequences between mGBP7 and hGBP1/mGBP2 and extend to their dimers. While hGBP1 and its close orthologue mGBP2 dimerize via their G domains, mGBP7 shows a variety of possible dimer structures, among them parallel and crossed-stalk conformations. The G domain is only partly involved in mGBP7 dimerization, which provides a rational why mGBP7, unlike hGBP1 and mGBP2, can dimerize in the absence of GTP. The different GBP dimer structures, which still exhibit hinge movements to certain degrees, are expected to encode diverging functions, such as a destabilization of pathogenic membranes or fusion of the parasitophorous vacuole membrane with the autophagic machinery.

## 1 Introduction

Dynamin superfamily proteins (DSPs) are mechanochemical enzymes, converting GTPase activity into some of the highest torques observed for proteins, which is then used for membrane fusion or fission.^1, 2^ In contrast to the smaller Rat sarcoma virus (Ras)-like GTPases, DSPs exhibit lower substrate affinity but higher basal hydrolysis rates, which is highly stimulated upon oligomerization.^3^ Furthermore, no accessory proteins, such as GTPase-activating proteins or guanine nucleotide exchange-factors are required since these functions are already encoded in the corresponding GTPase effector domain.^3, 4^ The oligomers of DSPs form ordered rings, lattices or helices, thereby tubulating membranes,^2^ and are thus involved in critical cellular functions, such as endocytosis, mitochondrial membrane tubulation, cell division, and vesiculation.^5, 6^

Since some of the DSPs have been studied separately, the nomenclature of domains is not unified, although they are functionally homologous and feature a modular composition of domains. Some examples are depicted in Fig. 1A. The GTPase domain (G domain) is the only conserved domain at the sequence level, and it shows the typical motifs of a nucleotide-binding domain (G1–G5, switch1 (SW1), switch2 (SW2)), in total spanning about 300 continuous residues. GTPase activity, although stimulated by multimerization, is usually not dependent upon it.^7, 8^ In the case of dynamin, an intramolecular interaction can replace intermolecular complementation of the G domains.^2, 9^ Other domains of DSPs are structurally conserved, even if they are not homologous in sequence. The bundle signaling element (BSE or neck or helical bundle 1) is an elongated helical domain that is in contact with the G domain via the so-called hinge 2 and connects to the stalk (or trunk or middle/effector-domain or helical bundle 2) domain via hinge 1. The membrane interaction can be mediated by a pleckstrin homology (PH) domain, the L4 loop, a variable domain (VD), a paddle or a C-terminal region (Fig. 1A and B).^2^

**Figure 1:**
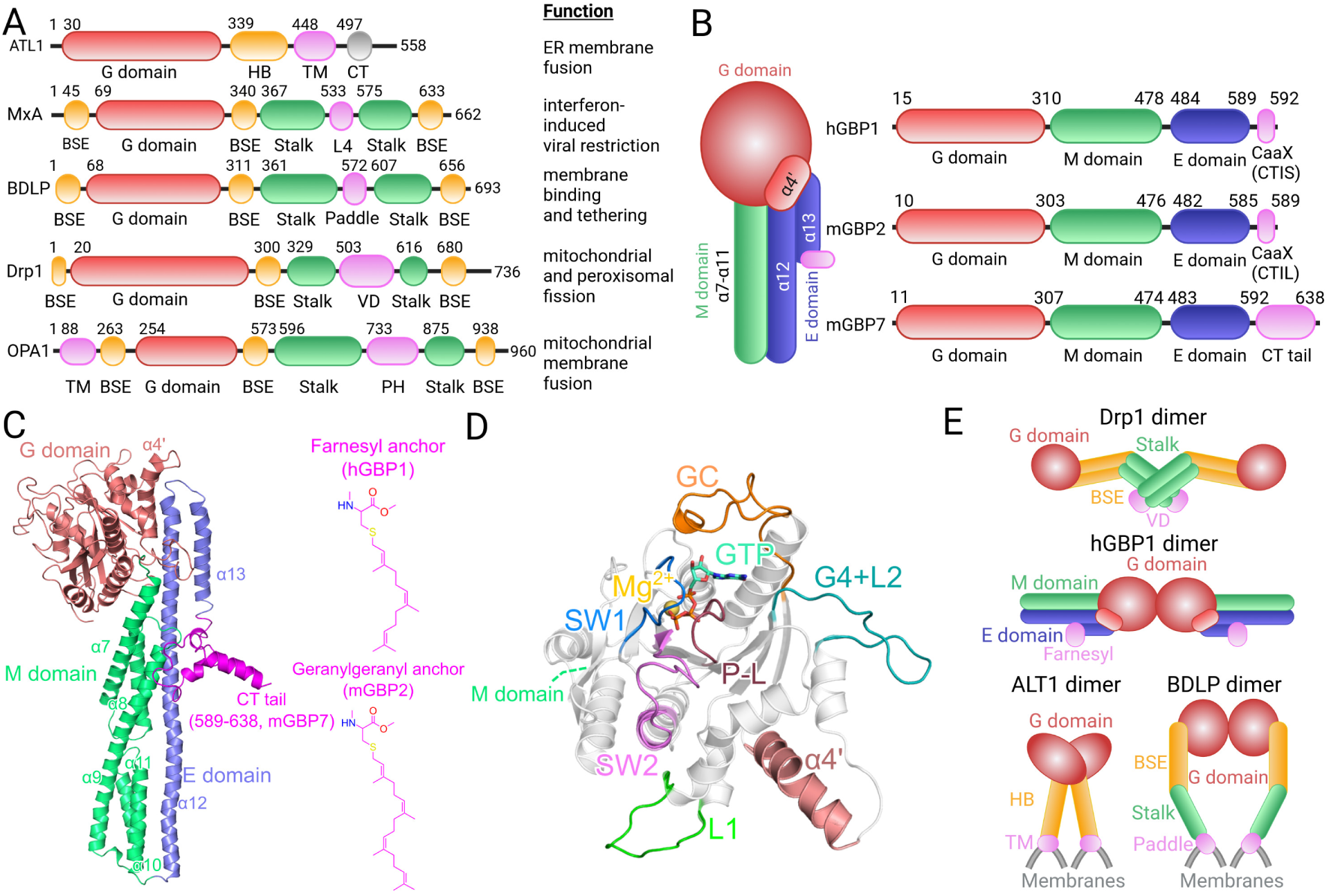
Structural organization and dimerization modes of DSPs and GBPs. **(A)** Schematic domain organization of selected DSPs: atlastin 1 (ATL1), myxovirus resistance protein 1 (MxA), bacterial dynamin like protein (BDLP), dynamin-related protein 1 (Drp1), OPA1 (from optic atrophy gene 1). ^2, 6, 10^ DSPs exhibit a highly conserved large GTPase domain (G domain, red), a structural element or domain for membrane binding (violet) which can be a transmembrane (TM) domain, the L4 loop of the G domain, a paddle domain, a variable (VD) domain, or a pleckstrin homology (PH) domain, two elongated *α*-helical bundle domains called stalk (green), and a bundle signaling element/helical bundle (BSE/HB, orange). The function of each DSPs is given on the right. **(B)** Schematic domain organization of GBPs. They are split into three domains: the large GTPase (G) domain (red), the middle (M) domain (green), and the effector (E) domain (blue), followed by a region responsible for membrane binding. It is either a CaaX box for isoprenylation or an elongated C-terminal (CT) tail. **(C)** The structure of mGBP7 is shown as cartoon. The helices discussed in this work (*α*4’, *α*7–13) are indicated. At the C-terminus, mGBP7 features a helical CT tail, while hGBP1 and mGBP2 become isoprenylated with either a farnesyl (hGBP1) or a geranylgeranyl (mGBP2) group (2D structures on the right). **(D)** Close-up of the G domain of mGBP7. The four conserved GTP-binding site motifs and other important structural elements are highlighted: P-L or G1 (dark red), SW1 or G2 (blue), SW2 or G3 (magenta), G4 plus loop L2 (G4+L2, turquoise), loop L1 (green), guanine cap (GC, red), and *α*4’ (red). The bound nucleotide GTP is shown with sticks and colored by atom type and Mg^2+^ is shown as a yellow sphere. **(E)** Known DSP dimerization modes are shown for Drp1 (PDB ID 4BEJ), ^11^ hGBP1 (PDB ID 2BC9) ^3^ featuring the G–G domain dimer, ^2^ ATL1 (PDB ID 3Q5D), ^12^ and BDLP (PDB ID 2W6D). ^13^ In general, the dimerization occurs via the stalk regions (Drp1) or via the G domains (ATL1, hGBP1, BDLP). Panels A, B, and E were created with BioRender.

One subfamily of DSPs is the guanylate binding protein family (GBPs, Fig. 1B), which are known, among others, for their anti-*Toxoplasma gondii*,^14–17^ anti-HIV,^18^ and anti-*Chlamydia trachomatis* ^19, 20^ activity. Using bioinformatic/phylogenetic methods, seven human GBPs (hGBP1 to hGBP7) and 11 murine GBPs (mGBP1 to mGBP11) have been identified.^15, 21^ In this work, we will concentrate on hGBP1, mGBP2, and mGBP7, continuing our previous studies of hGBP1,^22^ mGBP2,^23^ and mGBP7.^24^ The GBPs exhibit three domains (Fig. 1): (1) a GTPase domain (G domain) for GTP binding and hydrolysis (Fig. 1D), (2) a middle domain (M domain) for regulation, and (3) a GTP effector domain (E domain) for interactions with lipid membranes. The M and E domain (called M/E domain herein) give the GBPs an elongated shape. The E domain can contain a CaaX motif at the C-terminus, which is the case for hGBP1 and mGBP2, resulting in an isoprenyl lipid anchor after post-translational modification, while in the case of mGBP7 the protein harbors about 50 more residues at the C-terminus, which were shown to be essential for membrane binding of the protein (Fig. 1B and C).^24^ mGBP2 is the murine orthologue of the well-studied hGBP1. It has been shown that mGBP2 binds to the membrane of the *T. gondii* parasitophorous vacuole before mGBP7; however, the deletion of mGBP7 – compared to mGBP2 – results in a higher lethality of mice infected by the parasite.^16, 17^ The membrane of the parasitophorous vacuole originates from lipids of the host cell and the parasite hides within the vacuole to escape from the host cell defense response. GBPs cycle between monomeric, dimeric, and polymeric states, with the polymer being formed by thousands of subunits on the parasitophorous vacuole membrane of *T. gondii*.^25^

To achieve oligomerization, DSPs use up to four interfaces in the helical (stalk) region, and only once a full helical turn around the invaginated membrane is completed, the G domains come into contact with the membrane („grip“). The substrate GTP, which has previously been loaded, is now processed to GDP, which leads to a conformational change („pull“), where the dynamin filaments slide against each other, constricting the membrane. GDP is weaker in promoting oligomerization, possibly leading to dissociation once the cycle is complete.^26–28^ While nucleotides are necessary for oligomerization of most DSPs, mGBP7 can form dimers in the absence of GTP, whereas mGBP2 dimerization is mostly dependent on nucleotide binding.^24^ Nucleotide-independent oligomerization has also been found for OPA1 (from the optic atrophy gene 1, Fig. 1A).^10^ Moreover, GBPs are able to hydrolyze GDP further to GMP, which suggests that they remain in an oligomeric state even after hydrolysis. The second hydrolysis takes place either in a tetrameric state, or in a drastically extended conformation of the dimer.^29, 30^ According to Sistemich et al.,^31^ binding of GTP to hGBP1 leads to solvent exposure of the farnesyl anchor, then dimerization via the G domains takes place, and in the transition state, the helix *α*12 folds out and/or further oligomerization occurs. When GTP is scarce, these polymers dissociate, indicating that GMP cannot stabilize the polymer.^31^ Oligomers are also more strongly associated with membranes, due to the high local concentration of membrane-binding motifs.^32^ Several studies suggest functional heterodimerization as a key concept for DSPs,^33–35^ especially proteins that are localized on the same chromosome and are thus transcribed together tend to associate.^25, 36^ With regards to GBPs, this specifically applies to mGBP1/mGBP2 and mGBP3/mGBP7.

Membrane fission is likely to require several continuous cycles of constriction, until the membrane tubule thickness is thin enough for the two eventually separating bilayers to touch each other. Examples of DSPs enabling fission are dynamin, dynamin-related protein 1 (Drp1), and Dnm1p, a yest protein homologous to Drp1. Fusion, on the other hand, requires the association of at least two proteins to opposing membranes (tethering) and subsequent conformational change, putting them in close spatial proximity to initiate the fusion process.^27^ Fusion DSPs, such as atlastin, Sey1p, mitofusins, OPA1, MGM1p, and LeoA/B/C, have permanent membrane binding ability, function as dimers, and have a fused BSE/stalk domain.^27^ Intriguingly, some proteins can serve both functions, i.e., fission and fusion, such as Vps1p and NosBDLP.^37^ It is likely that different dimerization motifs could confer different functions. Dimer structures of DSPs that are deposited in the Protein Data Bank (PDB) display G domain dimers (hGBP1, Irga6, dynamin), crossed-stalk dimers (dynamin), parallel (BDLP, Sey1p) and antiparallel (SX9) stalk dimers (Fig. 1E), and higher-order oligomers that combine several of these interfaces.

Whether GBPs actively destabilize pathogenic membranes, including parasitophorous vacuole membranes, or fuse them within the autophagic machinery, or both, is still unknown. However, information about their dimerization patterns and dynamics could yield important hints. Thus far, only two partial dimer structures of GBPs are available.^3, 38^ The structure by Ghosh et al.^3^ is a dimer only formed by the G domains of hGBP1, while in the structure reported by Cui et al. two C-terminally truncated human hGBP5 molecules, comprising the G and M domains, interact lengthwise, with a cross formed at the base of the G domains. This dimerization mode is similar to the one known for atlastin, which cycles between this and the G domain dimer structure as seen for hGBP1.^39, 40^ Based on molecular dynamics (MD) simulations, we have previously unraveled a large-scale hinge motion taking place in monomeric and dimeric hGBP1^22^ and mGBP2,^23^ which is also present in the membrane-bound protein form. For the dimer structures, we assumed the presence of the G domain dimer as reported for hGBP1.^3^ Here, we test for the existence of alternative dimer structures of hGBP1, mGBP2, and mGBP7 by applying a combination of protein-protein docking, small-angle X-ray scattering (SAXS), and cross-linking mass spectrometry (XL-MS). The stability of the resulting dimer models is analyzed using MD simulations, which at the same time allows us to contrast the dynamics in the GBP monomers and dimers as well as to pinpoint the important residues in the dimerization interfaces. In this study, we provide evidence, that mGBP7 prefers different dimerization modes as compared to hGBP1 and mGBP2, which ties in with some differences in their sequences and dynamics that are reported here too.

## 2 Results

### 2.1 Sequence, structural and biochemical comparison of hGBP1, mGBP2 and mGBP7

First, we compare the protein sequences, molelular structures and biochemical properties of hGBP1, mGBP2, and mGBP7. Their most relevant properties are summarized in Table 1.

**Table 1:**
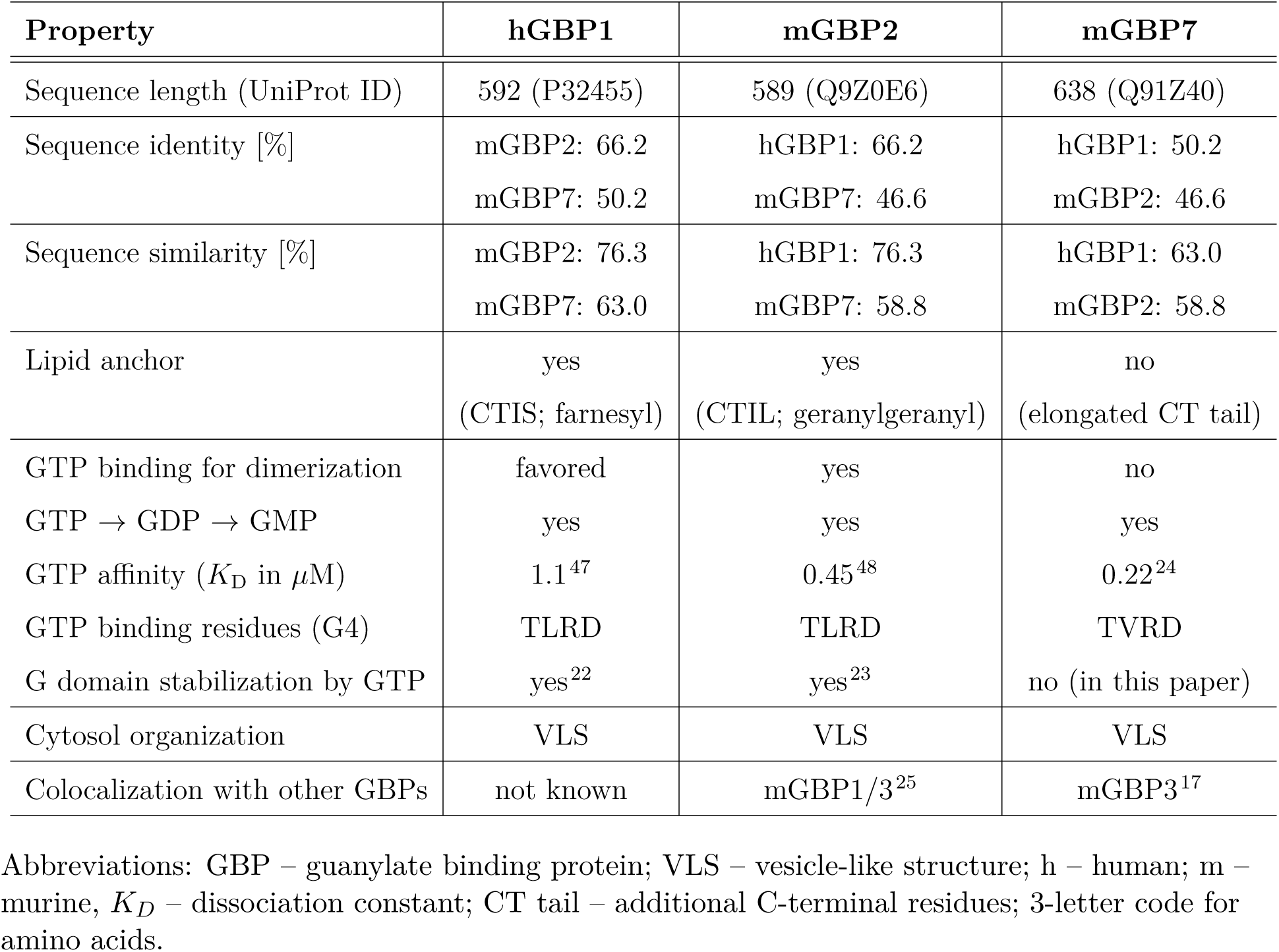
Summary of properties of hGBP1, mGBP2, and mGBP7. Abbreviations: GBP – guanylate binding protein; VLS – vesicle-like structure; h – human; m – murine, *K_D_* – dissociation constant; CT tail – additional C-terminal residues; 3-letter code for amino acids.

#### Membrane binding and protein assembly

All three GBPs are composed of a G, M, and E domain,^41^ with the M and E domain sometimes grouped together as an M/E domain (Fig. 1B and C). mGBP2 is a close orthologue of hGBP1 with 66.2% sequence identity and both feature a CaaX motif (residues 589–592) leading to isoprenylation. The mGBP7, on the other hand, contains 49 additional residues at the C-terminus (590–638, called CT tail) compared to mGBP2, which mediate the protein binding to a membrane (Fig. 1C).^24, 36^ Residues 598–620 were predicted as a transmembrane (TM) helical region using TMpred^42^ and mutation experiments indeed confirmed the CT tail to be essential for membrane binding of mGBP7.^24^ Thus, all three proteins can bind to lipid membranes. Moreover, in the cytosol, they form vesicle-like structures (VLS), which can also result from hetero-assembly of mGBP2 co-localizing with mGBP1/3 or mGBP7 with mGBP3.^17, 25, 31^ However, mGBP2 and mGBP7 do not co-localize in VLS.^25^ For hGBP1 it has been shown that in the presence of GTP and the farnesyl lipid anchor it polymerizes with the E domain being folded out, leading to the formation of hGBP1 disks that assemble into tubes and *in vitro*, they can tether membranes.^43–45^ Binding of GTP to hGBP1 shifts the monomer–dimer equilibrium in favor of the dimer, which enhances the GTP hydrolysis.^45^ However, this was suggested to loosen the contacts between the G domains defining the dimer, provoking E domain contacts to be formed, which is followed by GDP hydrolysis and dimer dissociation.^30, 32, 46^

#### GTP binding and hydrolysis

The G domain contains the four G motifs (G1–G4) needed for the GTP hydrolysis, the guanine cap, the loop 1 (L1), and the *α*4’ helix involved in membrane interactions (Fig. 1D). All three GBPs can hydrolyze GTP to GMP in a two-step mechanism, which is a big difference to other DSPs where the hydrolysis stops at GDP.^2, 49, 50^ However, none of the GBPs is able to directly bind GDP, i.e., it can only be processed as an intermediate following GTP hydrolysis.^24, 48, 49^ The GTP binding affinity is slightly different between the proteins, with mGBP7 having the highest and hGBP1 the lowest affinity (Table 1). To understand the differences in the GTP binding, we aligned the sequences of

hGBP1, mGBP2, and mGBP7 using T-coffee^51, 52^ and analyzed their amino acid compositions in terms of physicochemical properties (Fig. S1). This extends to a representation of their electrostatic potential surface (EPS) for the whole protein and the GTP binding site in particular (Fig. S2). All three GBPs are rather negatively charged on one side of the protein along the M/E domain, yet feature a mix of positive and negative charges on the opposite side, yet with a higher share of positive charges in the G domain on that protein side. Of the three proteins, mGBP7 is the most negative on either side, followed by mGBP2. However, the GTP binding region of mGBP2 is more positive than in hGBP1, which might explain mGBP2’s higher affinity for GTP.^47, 48^ Though this reasoning should be taken with caution, as the GTP affinity of mGBP7 is even higher, yet its GTP binding region is not more positive but contains mixed charges.

The G1 motif, also called phosphate-binding or P-loop (P-L), is highly conserved and is key for GTP binding and hydrolysis. The two most important residues are R48 serving as arginine finger and K51 being responsible for GTP hydrolysis. The K51A mutant drastically impairs GTP binding: in hGBP1 by *∼*50-fold (*K*_D_ = 53 *µ*M),^47^ in mGBP2 by *∼*100-fold (*K*_D_ = 44.1 *µ*M),^48^ and mGBP7 lost its GTP binding ability altogether.^24^ Moreover, only negligible GTP hydrolysis and no dimerization takes place for the K51A mutant of hGBP1 and mGBP2, while the mGBP7 mutant can still dimerize as it does not require GTP for this. The R48A mutant of hGBP1 showed a slightly increased GTP binding affinity (*K*_D_ = 0.34 *µ*M) but a decreased GTP hydrolysis, and dimerization still takes place.^48^ Kravets et al. further revealed in 2016 that the R48A mutant has a decreased capacity for recruitment to the parasitophorous vacuole membrane, while the K51A mutant lacks this ability completely, which is accompanied by its inability to polymerize/localize in a VLS and to control *T. gondii* growth at all.^25^ For hGBP1, Zhu et al. demonstrated in 2013 that the antiviral function against influenza virus A is reduced by the K51A mutant.^53^ These observations underscore the importance of the G1 motif for the function of these GBPs.

The sequences of the next two G motifs, G2 and G3 (also called switches SW1 and SW2), feature like G1 only minor changes among the three proteins. Only for the surrounding residues of SW1 we identified changes, such as Q72 and H74 in hGBP1 and mGBP2 became R72 and E74 in mGBP7. In hGBP1, the mutants Q72A, H74A, T75A and E99A led to a decreased GTP binding and hydrolysis with reduced GMP production.^47^ Another function of H74 was recently identified, as it was found to play a critical role as a distant molecular switch for the E domain releasing from the G domain of hGBP1.^44^ The SW2 of mGBP7 harbors more hydrophobic and also more positively charged residues, while in mGBP2 these are more polar and negatively charged than in hGBP1 and mGBP7. However, the key residues T75 of SW1 needed for interactions with Mg^2+^ as well as S73 of SW1 and E99 of SW2 required for coordinating the nucleophilic water molecule during the hydrolysis reaction are the same.^3, 47^ For mGBP2, only data of an E99A mutant is known, which shows decreased GTP hydrolysis, a low multimerization capability, and a reduced binding to parasitophorous vacuole membranes.^25, 48^ For mGBP7, there are no such mutation data available yet.

The G4 motif, which is responsible for mediating nucleotide specificity together with the loop 2 (G4+L2), is more positively charged in mGBP7 than in the other two GBPs. The first residues of that motif are not very different among the three proteins. An interesting change occurred at position 182 and 184 in mGBP2/7 and hGBP1, respectively, where a valine is found in mGBP7 as compared to a leucine in hGBP1 and mGBP2. The mutations D182N and D184N in hGBP1 and mGBP2, respectively, resulted in a 20 fold lower affinity for GTP and led to a reduction in the GTP hydrolysis rate, cooperativity, and GMP formation.^25, 47^ Moreover, the D182N mutant of mGBP2 might have a similar effect on multimerization and controlling *T. gondii* growth as the K51A mutant, which is dysfunctional with regard to GTP hydrolysis.^25, 48^ The position 182 is therefore an interesting candidate for future mutational studies of mGBP7 to assess its involvement in GTP binding and hydrolysis.

#### G domain loops

The most important element for dimerization of hGBP1 is the guanine cap, which is an unstructured loop region.^3, 22^ In hGBP1, two arginine residues, R240 and R244 exhibit a key role for the dimerization.^54^ Interestingly, mGBP7 has oppositely charged residues in that region. Examples are R244 in hGBP1 and R242 in mGBP2 that converted to D242 in mGBP7, and K246 in hGBP1 and Q244in mGBP2 became E244 in mGBP7 (Fig. S1). As a result, the overall charge of the guanine cap in mGBP7 is *−*3 while it is neutral in hGBP1 and mGBP2. The other larger loop region of the G domain is the loop 1 (L1), which in mGBP2 is more hydrophobic and polar but less charged than in hGBP1 and mGBP7. It can therefore interact with the geranylgeranyl lipid anchor and also the helix *α*12 of the E domain.^23^ The *α*4’ helix, which belongs to the G domain, also interacts with the *α*12 as well as with *α*13 via salt bridges, which keeps the E domain close to the G domain.^32^ For hGBP1 it was demonstrated that this “lock” can be broken by GTP hydrolysis, releasing the whole E domain from the G domain in order for hGBP1 to polymerize and tether membranes.^43, 44^ Interestingly, mutating the hGBP1 residues F171 and F175 in L1 and F229 in *α*4’ to alanine leads to a higher rate in GDP hydrolysis, even though these residues are quite far away from the active site.^44^ The same is true for the E227A/K228A mutant, which disrupts the intramolecular interactions between *α*4’ and *α*12/13, enabling release of the E domain from the G domain. Not much is known yet about the biochemical importance of the same residues in mGBP2 and mGBP7, yet their involvement in the protein dynamics and dimerization will be investigated within this study.

#### M/E domain

All three proteins harbor a considerable amount of charged residues in the M/E domain. A major difference is that the M domain of mGBP7 has an overall charge of +10, whereas that charge is *−*2 and 0 in hGBP1 and mGBP2, respectively (Fig. S1B). This might be of relevance for the membrane binding of mGBP7, especially since these positive charges are preferentially found on one side of the protein (Fig. S2A) and are augmented by an overall positive charge of +3 of the CT tail, which was already demonstrated to be essential for membrane interaction.^24^ Another common feature of the three protein sequences is that positively or negatively charged residues are repeated twice or more often and the opposite charges tend to cluster together. Examples are (i) triply repeated residues of the same charge, like ^417/419^EEE^419/421^ in mGBP2/hGBP1 and ^439^RKK^441^ in mGBP7; (ii) mixture of positively and negatively charged residues, often in repeats, like ^389^EKKRDD^394^ in hGBP1, ^389^KRD^391^ in mGBP2, and ^388^EEKRED^393^ in mGBP7. The second variant occurs more often in all three GBPs. In mGBP7, however, there are more breaks by polar and hydrophobic residues between the charged stretches, which are therefore more distributed along the helices. The helices *α*12/13 are not amphipathic in either of the three GBPs, as the charged residues are not only solvent-exposed, some of them are also buried (Fig. S2B). For hGBP1 we showed that some of the charged residues of *α*12 form salt bridges with the M domain that are crucial for the stability of that long helix.^22^ The *α*13 helices of hGBP1 and mGBP7 feature clusters of positive charges, ^582^KMRRRK^587^ and ^567^KRK^569^, respectively. Alanine-scanning mutagenesis of the three arginines in *α*13 helix of hGBP1 ablated the farnesylated protein’s ability to bind to bacterial outer membranes.^55^ The mGBP2 does not have repeated lysine or arginine residues in *α*13 directly before the CaaX motif, only a single lysine residue (K585), which one can expect to help in membrane binding together with the longer geranylgeranyl lipid anchor compared to the farnesyl group in hGBP1. The sequence ^567^KRK^569^ in mGBP7 is directly before the CT tail, which was demonstrated to be essential for membrane binding.^24^ The CT tail has a +3 charge, coming from six positively and three negatively charged residues. Of the other 40 CT tail residues, there are 27 aromatic/hydrophobic ones. This combination of charges and hydrophobicity allows the CT tail to take over the role of a membrane anchor, replacing the function of the isoprenyl group of hGBP1 and mGBP2.

### 2.2 Molecular dynamics of monomeric GBPs

Before establishing dimer models of the three GBPs under investigation, we compare the dynamics of their monomers, on the one hand to further elucidate possible differences between hGBP1, mGBP2, and mGBP7, and on the other hand to have the monomer dynamics as a reference for the dynamics of the proteins when being oligomerized. The dynamics of hGBP1 and mGBP2 was discussed in detail in our previous works,^22, 23^ while the dynamics of mGBP7 is being unraveled in this study. In all three cases, Hamiltonian replica exchange molecular dynamics (HREMD) simulations were applied to the monomeric proteins. All three proteins were simulated in their apo-state (denoted hGBP1_apo_, mGBP2_apo_, and mGBP7_apo_ here) and for mGBP2 and mGBP7 we also considered the GTP-bound state (mGBP2_GTP_, mGBP7_GTP_). For hGBP1 and mGBP7 we used 30 *×* 400 ns MD replicas and for mGBP2 it was 16 *×* 200 ns MD replicas (see Table 2 for a list of all simulations). For mGBP2 we had shown that the smaller HREMD setup was sufficient as no further protein states were sampled in a 40 *×* 400 ns HREMD simulation.^23^ For the analysis, we considered the protein conformations collected in the HREMD target replica that had no modifications applied to the potential energy function of either hGBP1, mGBP2, or mGBP7.

**Table 2:** Summary of simulations included in this work.*^a^*.

| Simulation | System | Size in atoms | Runs | Length | Cumulated time |
| --- | --- | --- | --- | --- | --- |
| HREMD<sup>b</sup> | hGBP1<sub>apo</sub> | 335,553 | 1 × 30 | 400 ns | 12 μs |
| HREMD<sup>c</sup> | mGBP2<sub>apo</sub> | 131,493 | 1 × 16 | 200 ns | 3.2 μs |
| HREMD<sup>c</sup> | mGBP2<sub>GTP</sub> | 129,116 | 1 × 16 | 200 ns | 3.2 μs |
| HREMD | mGBP7<sub>apo</sub> | 178,800 | 1 × 30 | 400 ns | 12 μs |
| HREMD | mGBP7<sub>GTP</sub> | 178,019 | 1 × 30 | 400 ns | 12 μs |
| MD | hGBP1 dimer | 556,008 | 1 | 100 ns | 100 ns |
| MD | mGBP2 dimer | 556,729 | 1 | 100 ns | 100 ns |
| MD | mGBP7 dimer, model 1 | 538,254 | 1 | 100 ns | 100 ns |
| MD | mGBP7 dimer, model 2 | 308,874 | 1 | 100 ns | 100 ns |
| MD | mGBP7 dimer, model 3 | 308,763 | 1 | 100 ns | 100 ns |
| MD | mGBP7 dimer, model 4 | 308,514 | 1 | 100 ns | 100 ns |
| MD | mGBP7 dimer, model 5 | 308,763 | 1 | 100 ns | 100 ns |
| **Total simulation time** |  |  |  |  | **36.7 μs** |

<sup>a</sup> The simulations reported in this work were either performed in the supercomputer JURECA at the Jülich Supercomputing Centre,<sup>84</sup> in SuperMUC-NG at the Leibniz-Rechenzentrum (LRZ) in Munich, or in HILBERT at Heinrich Heine University Düsseldorf.
<sup>b</sup> This trajectory originates from reference<sup>22</sup>.
<sup>c</sup> This trajectory originates from reference<sup>23</sup>.

#### Overall protein flexibility

The flexibility of the protein monomers was first quantified by the root mean square fluctuations (RMSF) of the C*_α_* atoms (Fig. 2 and S3). Among the three proteins, hGBP1_apo_ is the most flexible with RMSF values of up to 15.8 Å in the M/E domain, followed by mGBP2 with maximal RMSF values of 5.1 Å at residue 481, which is at the tip of the M/E domain, in the apo-state and 6.4 Å at residue 422 of *α*10 in the GTP-bound state, while mGBP7 is the least flexible with maximal RMSF values at residue 588 of *α*13, reaching 4.1 Å in mGBP7_apo_ and 4.9 Å in mGBP7_GTP_. The different protein flexibilities are also visible in Figure 2 where the protein structures are colored according to the RMSF values.

**Figure 2:**
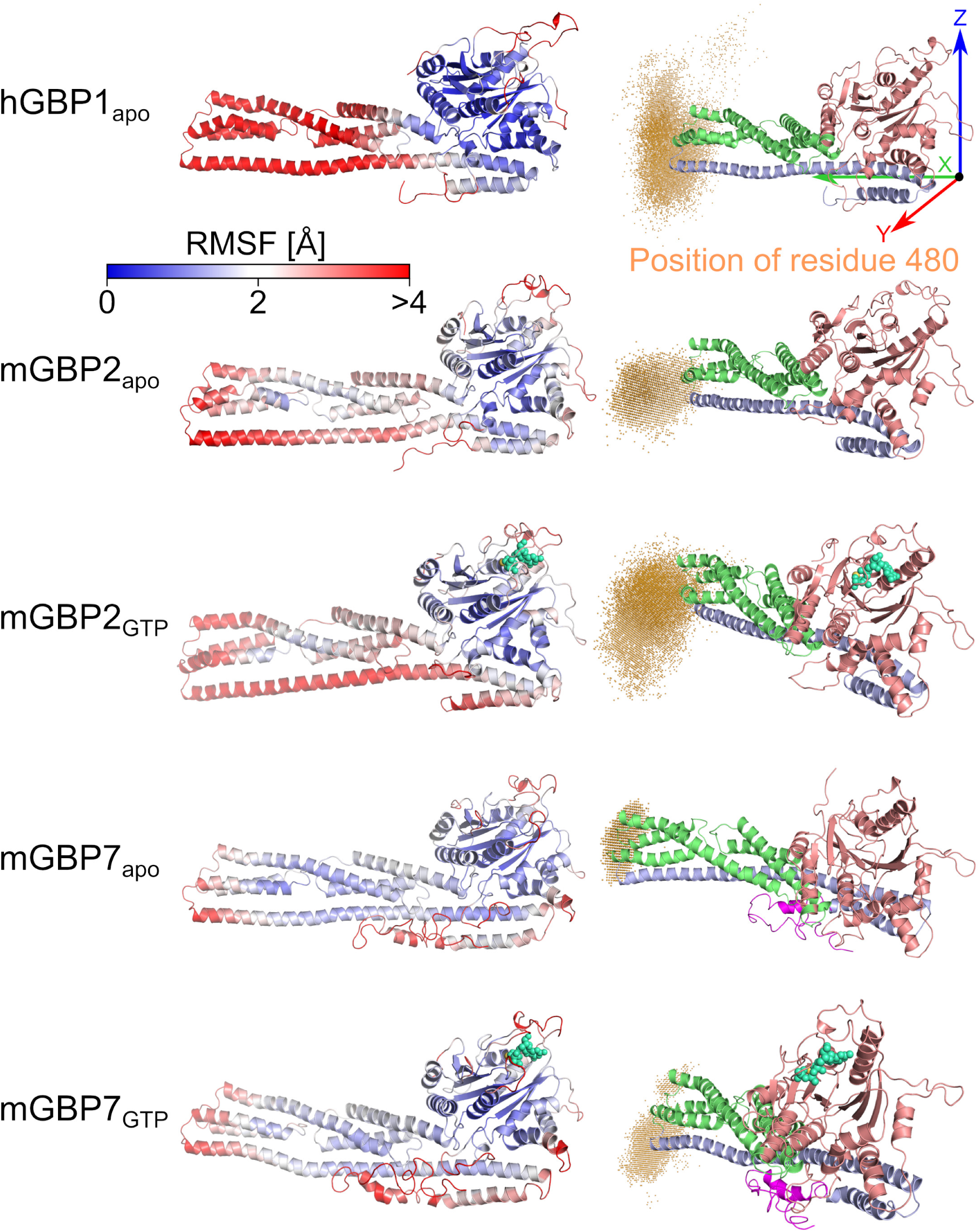
Structural fluctuations of the monomeric hGBP1, mGBP2, and mGBP7. **(Left)** The protein flexibility, as quantified by the RMSF of the C*_α_* atoms during the respective HREMD simulation, is projected onto the initial structure of hGBP1_apo_, mGBP2_apo_, mGBP2_GTP_, mGBP7_apo_ and mGBP7_GTP_. Rigid residues are colored in blue and flexible residues are shown in red, according to the color scale at the top. **(Right)** The spatial distribution of the residue 480 (i.e., the tip of the M/E domain) is shown as orange cloud to illustrate the hinge motion of the corresponding GBP monomer. Taking hGBP1_apo_ as an example, the definition of the coordinate system is shown for the quantification of the hinge motion using the motions of the residue 480 with respect to the initial structure, as measured by Δ*x,* Δ*y,* Δ*z* and *d*_480_.

#### G domain motions

In mGBP2, binding of GTP stabilizes the G motifs and loops of the G domain, especially also the guanine cap, which is not so much the case for mGBP7. Except for the P-L motif, these motifs and loops in mGBP7_GTP_ remain flexible with RMSF values above 2 Å (Figs. 2 and S3).

To further characterize the motions of the G domain motifs and loops, we performed a clustering analysis of the G domains, with the simulation snapshots being fitted onto the *β*-sheets of the corresponding G domain. The resulting numbers of clusters, their populations, and maximal structural differences between the clusters are summarized in Table S1. The cluster data show that among the apo-states, mGBP2 has the most stable G domain, which becomes even stiffer in mGBP2_GTP_, except for L1, which shows only an insignificant RMSF decrease, and *α*4’, whose flexibility even increased slightly (3 instead 2 clusters). In mGBP7, the G motifs and loops are generally more flexible than in mGBP2 and the stabilization by GTP bindings is also smaller. This is most significant for the guanine cap, for which the number of clusters decreases only by 14% from mGBP7_apo_ to mGBP7_GTP_ as compared to a 90% reduction in mGBP2. The other structural G domain elements of mGBP7 experience larger decreases in the number of clusters, yet apart from P-L and G4+L2 they remain rather flexible in mGBP7_GTP_. This can also be seen in the structural presentation of the clusters in Fig. S4. The different behavior in these mGBP7 motifs and loops as compared to hGBP1 and mGBP2 largely correlates with the amount and type of changes in their amino acid composition. In particular the negative charge of the guanine cap in mGBP7, as opposed to charge neutrality of that region in the other two proteins, might cause its generally higher flexibility and electrostatic repulsion from GTP. This in turn prevents its stabilization in mGBP7_GTP_ as seen in mGBP2_GTP_ where the guanine cap becomes more ordered (Fig. S4). Interestingly, the relatively high flexibility of the G domain of mGBP7 does not affect the binding affinity of GTP, which is even somewhat higher for mGBP7 than for hGBP1 and mGBP2 (Table 1). It thus seems to be sufficient if the amino acid residues directly involved in GTP binding, such as R48 and K51 become rigid if GTP is present in order to enable a stable GTP binding site. It is interesting to note that mGBP7 shows a higher dynamics in the G domain independent of the GTP loading state, while its M/E domain is more rigid than in the other two GBPs.

#### Hinge motion of the M/E domain

The dynamics of the M/E domain is characterized as a hinge motion, as first discovered for hGBP1^22^ and then confirmed for mGBP2.^23^ To quantify that motion, we computed the motions of residue 480 at the tip of the M/E domain relative to its position in the crystal structure or homology model during the HREMD simulations. These motions of residue 480 are provided as changes in its Cartesian coordinates (Δ*x*, Δ*y*, Δ*z*, see Fig. 2 top right for the definition of the coordinate system) as well as its Euclidean distance between the current and reference position (*d*_480_). The resulting data are plotted as statistical box plots in Figs. S5 and S6, and are illustrated by the spatial distribution of residue 480 (Fig. 2). The difference in the three GBPs is mainly described by Δ*y*, while Δ*x* and Δ*z* are similar. For hGBP1, motions in all directions are possible, but motions in +Δ*y* and *−*Δ*z* are favored and resemble a jack-knife where the tip of the M/E domain is moving towards the G domain. In hGBP1, the absolute motion of residue 480 is the highest of all three GBPs, with the *d*_480_ values reaching up 69.8 Å. The tip of the M/E domain of mGBP2 also favors motions into +Δ*y* direction, in both its apo and GTP-bound state, which is accompanied by *−*Δ*x* motions as a result of the jack-knifing of the domain. With regard to motions along the *z* coordinate, we see a displacement towards *−*Δ*z* in mGBP2_apo_, while with GTP both directions of Δ*z* are equally achieved. We revealed that the motions in the +Δ*z* direction are accompanied by a stabilization of the salt bridges between *α*4’ and *α*12/13, which became possible due to a displacement of *α*4’ towards the E domain as a result of GTP binding.^23^ The maximal motion of residue 480 in mGBP2 is slightly lower than in hGBP1, but can also reach values of about 60 Å (mGBP2_apo_: max. *d*_480_ = 62.4 Å; mGBP2_GTP_: max. *d*_480_ = 58.9 Å). In mGBP7_apo_, the motion into *−*Δ*y* and +Δ*z* direction is preferred, while in mGBP7_GTP_ the M/E domain tip chooses to move into the opposite direction, i.e., into +Δ*y* and *−*Δ*z* direction. This implies that, as in mGBP2, GTP binding affects the direction of the hinge motion, yet in the opposite way, which correlates with the different structural effects that GTP binding has on the G domains of mGBP7 and mGBP2. Another difference is that Δ*x* is almost zero in mGBP7. This results from the overall smaller jack-knifing in mGBP7, as revealed by the maximal displacement of residue 480 (mGBP7_apo_: max. *d*_480_ = 35.2 Å; mGBP7_GTP_: max. *d*_480_ = 37.5 Å). This can be also seen from the spatial distribution of that residue in Fig. 2.

#### Salt bridges

We finish the analysis of the monomeric GBP simulations by assessing the stability of intramolecular salt bridges that are thought to be relevant during protein dimerization, in particular salt bridges within the guanine cap and between *α*4’and *α*12/13. The salt bridges in question are shown in Fig. 3A and their occurrences during the simulations are summarized in Fig. 3B. In hGBP1_apo_, the salt bridge between K63 and E256 or E257 in the guanine cap has an occurrence below 15%, while a very stable salt bridge with 98% probability formed between R227 of *α*4’ and E556 or E558 of *α*12. In the corresponding simulation of mGBP2_apo_, the salt bridge in the guanine cap (R62–E253) was more stable, with a population of 40%. However, upon GTP binding this probability decreased to 16%, which can be explained by the ordering of the guanine cap in mGBP2_GTP_, allowing the residue R62 to also interact with E249 with 77% probability (only 1.7% in mGBP2_apo_). The salt bridges K226–E573 and R231–E554 tethering *α*12/13 to *α*4’, are stable in both mGBP2 states with populations of over 70%, while the salt bridge R225–E561 does not exist in either monomer state. In mGBP7, the salt bridge R57–E254 in the guanine cap is stable independent of the GTP binding state, which agrees to the observation that also the structure of the mGBP7 guanine cap is hardly affected by GTP binding. The other possible salt bridge, R57–D255, is only formed in mGBP7_GTP_, yet also only with 9% probability. With regard to the *α*4’–*α*12/13 interactions in mGBP7, there are several possibilities for salt bridges, in particular between E222 and R566/K567 as well as R225 and D558/E562. They were all stable in both mGBP7_apo_ and mGBP7_GTP_ with occupancies above 50%. The only exception is the E222–R566 salt bridge in mGBP7_apo_ for which only a 13% occupancy was found.

**Figure 3:**
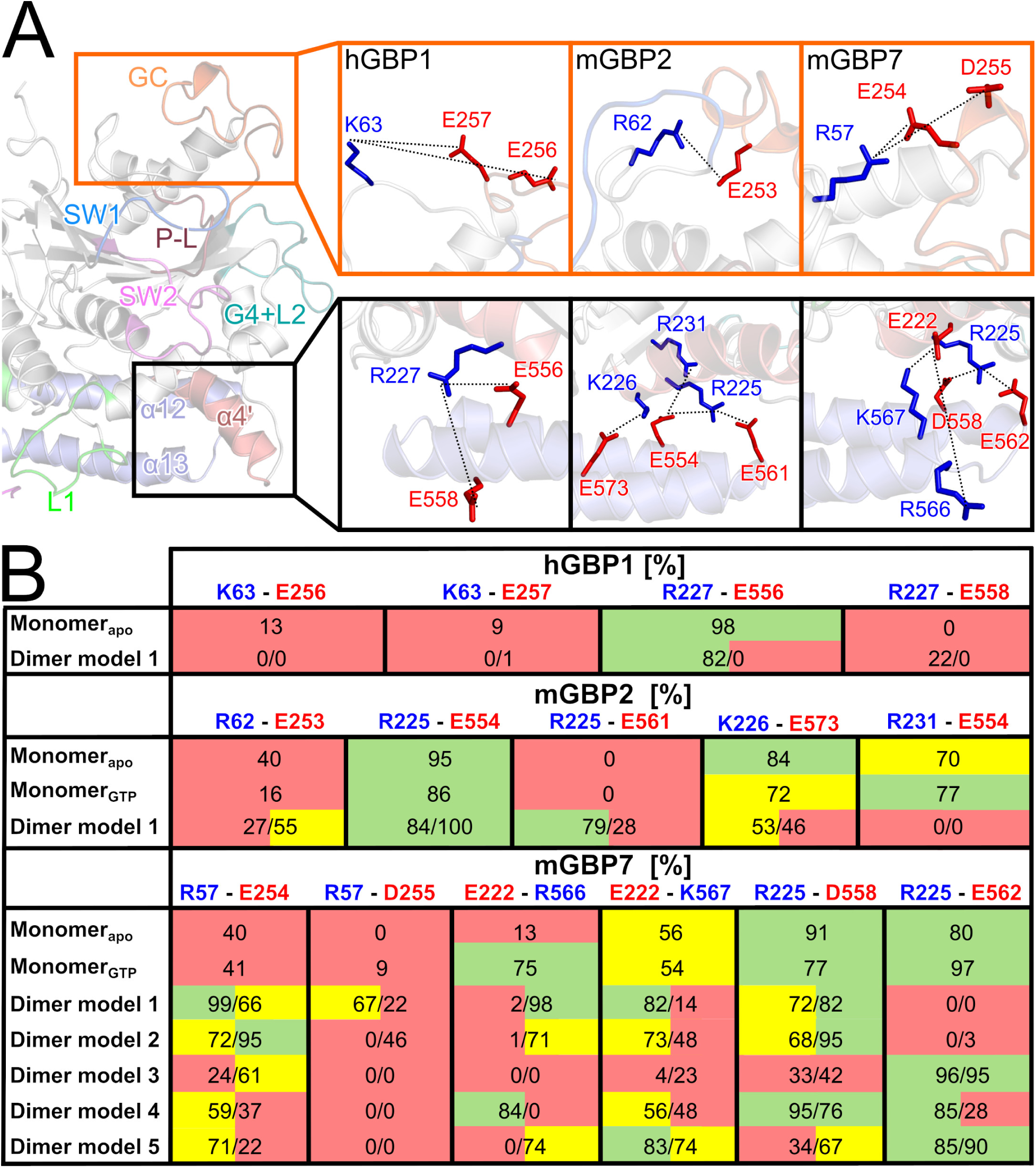
Intramolecular salt bridges relevant for dimerization. **(A)** On the left, parts of the G and the E domain are shown. The location of the salt bridges within the guanine cap and between *α*4’ and *α*12/13 are indicated by an orange and a black box, respectively. On the right, the zoomed views of these areas are shown for hGBP1, mGBP2, and mGBP7. The possible salt bridges are indicated by dotted lines between the residues being involved, which are labeled and their side chains highlighted as red and blue sticks for negatively and positively charged residues, respectively. **(B)** Occupancy (in %) of the salt bridges during the simulations. The results for chain 1 and chain 2 are given separately for the dimers. The results are colored based on the occupancy: 0–49%, red; 50–74%, yellow; 75–100%, green.

An interesting difference between mGBP7 and the other two GBPs is that *α*4’ of mGBP7 donates a positively and a negatively charged residue for the formation of salt bridges with *α*12/13, whereas in hGBP1 and mGBP2 only positively charged residues in *α*4’ are available for electrostatic interactions with *α*12/13. Apart from that, the overall conclusion is that in all three proteins there are stable salt bridges between the G and the E domain that kept the three helices *α*4’, *α*12 and *α*13 closely together independent of the GTP loading state.

### 2.3 Dimer models of GBPs

Our long-term goal is to elucidate the mode of action of membrane-bound GBP multimers involving up to thousands of subunits. To reach that goal, we first need structures for the polymerized GBPs and an understanding of their dynamics, especially also in comparison to the dynamics of their smallest organizational subunits, the monomers, which we just presented. Here, we continue by providing possible structure models for the smallest oligomer, the dimer, and assess the stability and motions of these GBP dimers.

#### 2.3.1 Model generation

We created structural models for the dimers of hGBP1, mGBP2, and mGBP7 based on experimental data, taken from the literature if available or produced within this study, combined with protein–protein docking. In the case of hGBP1, we constructed the dimer model using the crystal structure that exists for the dimer of its G domain (PDB IDs: 2B8W, 2B92 and 2BC9^3^) and aligning the G domains of two hGBP1 molecules with that structure (Fig. 4). It should be noted that we modeled the hGBP1 and mGBP2 dimers with GTP being bound as both proteins only notably dimerize in the presence of GTP,^3, 41, 48^ whereas the mGBP7 was considered in its apo-state as the dimerization of this protein is not affected by GTP^24^ and the experimental mGBP7 data obtained here was also recorded without GTP. We tested for the existence of alternative hGBP1 dimerization motifs using ClusPro,^56^ which is a physics-based protein-protein docking program. As such, it does not incorporate structural information from the PDB.^57^ Nonetheless, the best hGBP1 dimer structure proposed by ClusPro is very similar to the hGBP1 dimer created from the structure of the G domain dimer. Their RMSD from each other is only 5.0 Å, which is small for the system size under consideration, and their topological similarity based on the so-called TM-score calculated with MMalign^58^ is rather high with a value of 0.73. The TM-score can range between 0 and 1, with 1 corresponding to a perfect match between two structures, a TM-score of *>* 0.5 indicating that two structures have a similar topology, and a TM-score *<* 0.17 suggesting that the structural similarity is close to random . The ClusPro docking scores for the other hGBP1 dimer models were low, and we therefore did not consider them in our further study. A similar picture emerged for mGBP2 as a result of its 76.3% sequence similarity with hGBP1. The most likely mode of dimerization predicted by ClusPro is via the G domains, resulting in an mGBP2 dimer model very similar to the crystal structure-based one for hGBP1 (Fig. 4). The following five best mGBP2 dimer models are also all G domain dimers, with different rotations of the two proteins with respect to each other, suggesting that this interface is quite robust. However, these models had considerably lower docking scores than the top model. The application of AlphaFold-Multimer^59^ to mGBP2 produced the same outcome, i.e., only G domain dimers were suggested.^23^ Therefore, only the top mGBP2 dimer model produced by ClusPro was considered here.

**Figure 4:**
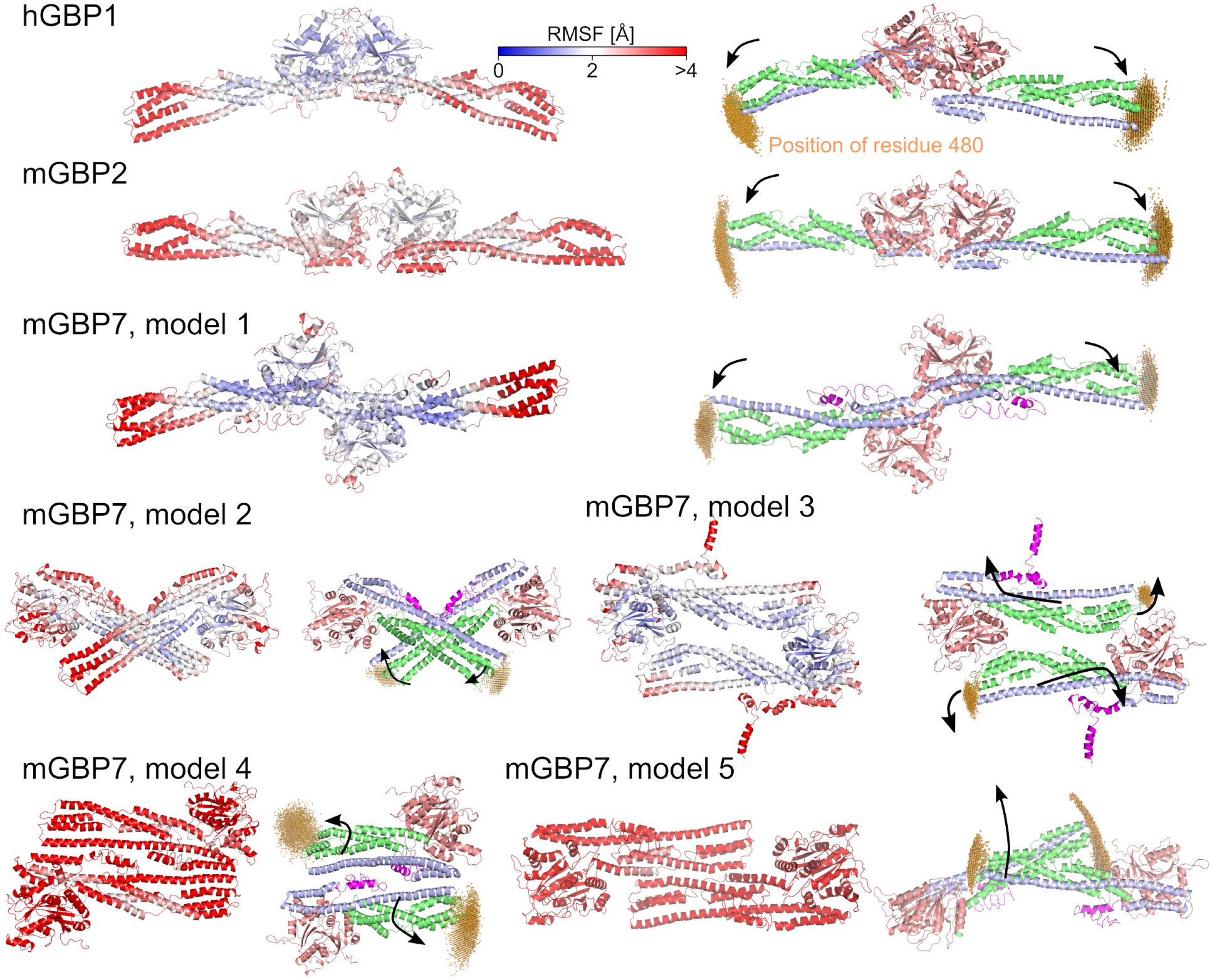
Structural fluctuations of the hGBP1, mGBP2, and mGBP7 dimers. The dimer models considered in this work are shown. The overall stability of the dimers was quantified by the RMSF of the C*_α_* atoms during the MD simulations of these dimers. The RMSF was calculated by aligning the whole dimer and is projected onto the initial dimer model, where rigid residues are colored in blue and flexible residues in red (color scale at the top). The degree of the hinge motion of the M/E domains was determined by the spatial distributions of the residue 480, which are shown as orange clouds. The coordinates used for measuring the hinge motion is the same as in Fig 2, with each chain of the dimers being aligned individually in that coordinate system. The black arrows indicate the main motions resulting from PCA analysis.

For mGBP7, the situation is different. First, ClusPro predicts three dimerization modes of equal likelihood for mGBP7. While one of them agrees nicely to the XL-MS data gathered here (discussed below), none of them could be satisfactorily fit into the molecular envelope reconstructed from our SAXS data. Therefore, we generated two further mGBP7 dimer models by fitting two mGBP7 proteins into molecular envelopes corresponding to the SAXS data using two different fitting techniques (discussed below). In total, we thus produced five mGBP7 dimer models, of which models 1–3 are from the ClusPro prediction and models 4 and 5 are from the SAXS data fitting (Fig 4). None of the mGBP7 dimers considered here involves the G domain interface seen for hGBP1 and predicted for mGBP2. ClusPro did not produce such a dimer model. Interestingly, the application of AlphaFold-Multimer^59^ to mGBP7 creates only G domain dimers (Fig. S7), yet their DockQ scores are low with respect to mGBP2 as reference. This indicates a poor match of amino acids at the dimer interface,^60^ which can be rationalized with the different amino acid compositions of the guanine caps of both proteins. Since the G domain dimer model is also not in agreement with of our experimental data for mGBP7 dimers, we did not consider it further.

In the following, the G domain dimer of hGBP1 and mGBP2 and dimer models 1–5 of mGBP7 are discussed in detail. To assess the stability and flexibility of the dimers, we performed 100 ns MD simulations per dimer and calculated the RMSF and stalk tip motions. Moreover, a principal component analysis (PCA) was applied to the MD data to identify the most prominent collective motions. To assess the dimer interface, we calculated the dimer contact area, analyzed selected salt bridges (see Fig. 3A), and determined the energies of the intermolecular contacts.

#### 2.3.2 Structural details of the hGBP1 and mGBP2 dimers

The dimerization via the G domains as considered for hGBP1 and mGBP2 allows to bury a large protein area. Of all dimer models discussed here, the hGBP1 dimer has the largest interface, with 5,314 Å^2^, and also the mGBP2 dimer model is stabilized by a large interaction area of 3,129 Å^2^.

To assess the dimer stability and flexibilities, we make use of the RMSF again. When aligning the whole dimer for the RMSF calculation, we obtain a measure for the overall dimer stability (Fig. 4), which reveals dimer model for hGBP1 to be more rigid than for mGBP2. The flexibilities of the chains within the dimers (Fig. S8), calculated after alignment of each chain, are very similar to the RMSF profiles of the corresponding monomers, suggesting that dimerization does not have a large effect on protein movements. As for the monomers, the stalk tips are the most flexible, with RMSF values of up to 8 Å. Notably, in the monomer state mGBP2 is less flexible than hGBP1, while it is the other way round for the dimers. Other flexible regions in bother dimers are L1, G4+L2, *α*4’, and the guanine cap. Nonetheless, compared to the monomers, the guanine cap as well as SW1 are stabilized, which correlates with salt bridge formation between the two chains (Fig. 3B). The two hinge motions per dimer, as described by the spatial distribution of residue 480 (Fig. 4) and the motions of this residue relative to its starting structure as defined by the Δ*x, y, z* values (Fig. S5) are similar, but they are more pronounced in mGBP2 as evidenced by higher *d*_480_ values due to larger motions along almost all axes. The preferred directions of the motions appear along +Δ*y/ ±* Δ*z*. The two hinge motions in either dimer are correlated with each other, as revealed by the PCA. The first principal motion can be likened to a butterfly motion (see arrows in Fig. 4), leading to dimer geometries that are more curved or C-shaped, which is accompanied by the formation of an intramolecular contact between residues K62 and D255 (Fig. 3) and unfurling of the C-terminal regions (yet more so in one of the two chains per dimer). The motion requires a certain structural flexibility at the dimer interface, especially in all GTPase motifs. The second principal motion is comparable to the first one, yet involves more screwing of the M/E domain in the lateral dimension.

The structure of the guanine cap is stabilized in both dimers compared to the monomeric proteins, yet the stabilization originates from partly different interactions in the hGBP1 and mGBP2 dimer (Fig. 3). The occurrence of the intramolecular salt bridge K63–E256/E257, which already in the monomer only occurred with a probability of *∼*10%, is abolished in the hGBP1 dimer. The corresponding salt bridge R62–E253 in the mGBP2 dimer gained in strength with respect to the mGBP2_GTP_ monomer, reaching a chain-averaged occurrence similar to the 40% observed for the mGBP2_apo_ monomer. The *α*4’–*α*12/13 lock is overall weakened in both dimers, but more so in the hGBP1 dimer. In mGBP2, where more salt bridges between *α*4’ and *α*12/13 are possible, the R225–E554 interaction is of particular stability and remains intact with a *>* 84% probability in all mGBP2 systems studied here. With regard to intermolecular interactions, the hGBP1 dimer harbors strong (about *−*80 kJ/mol) contacts involving both guanine caps, such as E261–K252’, K252–D239’, and R245–E251’/E256’ as well as E105–K209’ (where the prime indicates that these residues belong to the other chain), which are mostly Coulombic in nature. The mGBP2 dimer is also held together by interactions between both guanine caps, but further involves interactions of residues in the 130ies and of the G4+L2 motif that form contacts with their respective counterpart in the other chain. The single strongest interaction is that of D237–R238’, with an interaction energy of *−*120 kJ/mol (Fig. S9).

#### 2.3.3 Experimental data for mGBP7 dimers

##### XL-MS

The aim of XL-MS was to obtain information on the mGBP7 dimerization interface. To this end, recombinant mGBP7 was incubated in solution with either of two crosslinkers: bis(sulfosuccinimidyl) suberate (BS3, 11.4 Å) or disuccinimidyl sulfoxide (DSSO, 10.1 Å). Both crosslinkers react mainly with lysine residues and can link two lysines within a distance of 12 Å. Potential mGBP7 dimers and monomers were separated in a polyacrylamide gel and separately analyzed by mass spectrometry. Crosslinked peptide pairs were subsequently mapped to the different mGBP7 dimer models created with ClusPro and to the mGBP7 monomer (summarized in Table S2). A focus of the analysis was the identification of potential intermolecular contact sites within the homodimer, for which especially bridges between the same residues (like 216–216’, 565–565’, 567–567’) – as revealed by BS3 crosslinking – were informative. Of the three mGBP7 dimer models that resulted from docking, only model 1, which has an elongated shape with the dimer interface being formed via the two G domains (Fig. 4), agrees with our crosslink information (Fig. 5A and B), while the G domain dimers predicted by AlphaFold disagree. The XL-MS intermolecular contacts can be mapped to the *α*4’ of the G domain and *α*13 of the E domain. Moreover, the interactions 205–567’, 216–576’ and 567–576’ for BS3 as well as 106–216’, 205–554’, 205–565’ and 557–567’ for DSSO are in agreement with the intermolecular contacts in this model. In conclusion, this is the most probable model that fits to the XL-MS results.

**Figure 5:**
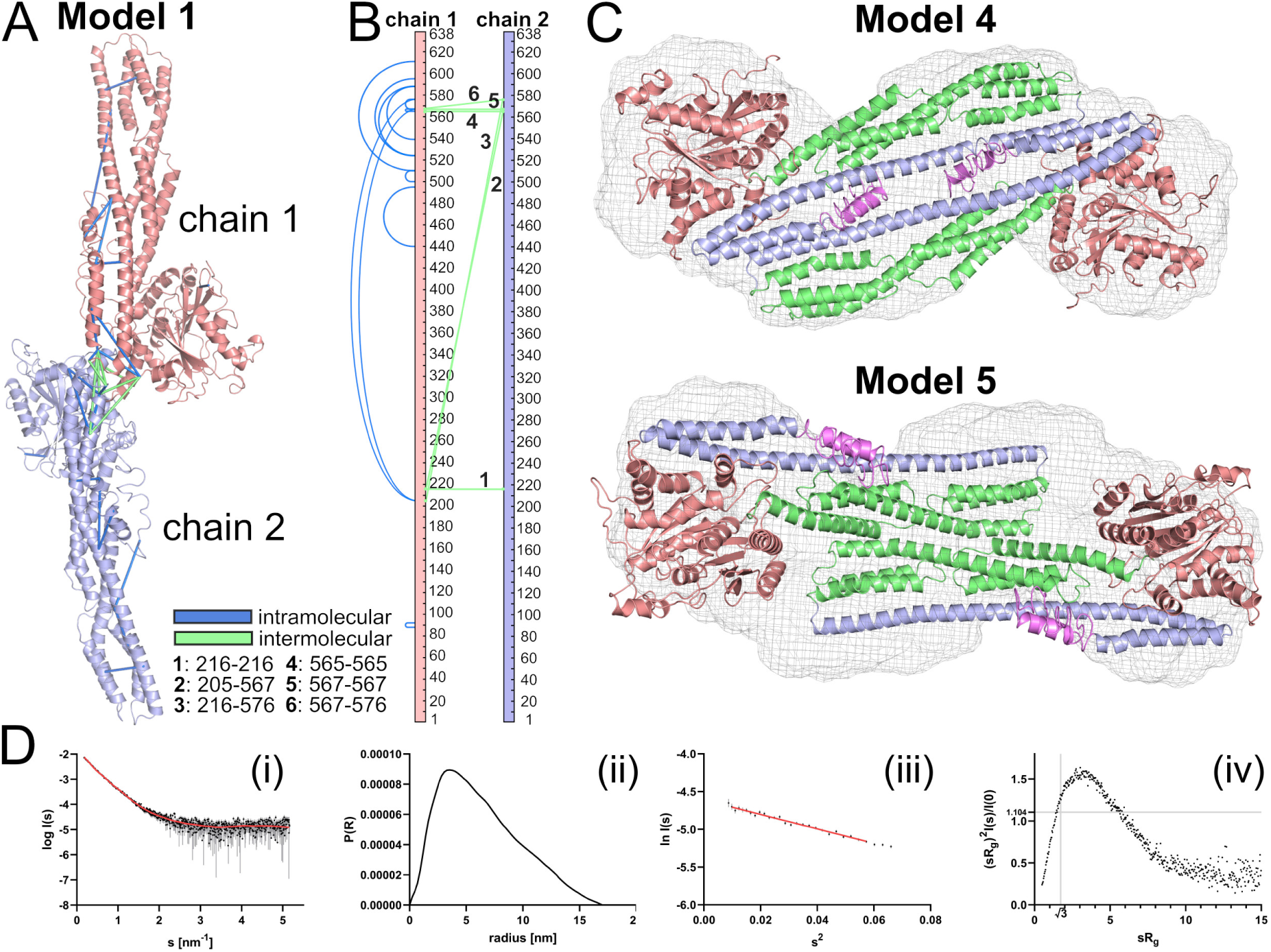
Results of the XL-MS and SAXS data of mGBP7 dimer. **(A,B)** Visualization of the crosslinks matching model 1 for the mGBP7 dimer. Mapped intermolecular crosslinks (numbers of the linked residues are given) are shown in green, while intramolecular crosslinks are colored in blue. Crosslinks are indicated in the 3D model in (A) and in a linear sequence plot (created with xiNET ^61^) in (B). **(C)** Calculated SAXS envelope for mGBP7 with dimer models fit into it: (top) model 4 created with GASBOR ^62^ and (bottom) model 5 created with SASREF. ^63^ **(D)** (i) SAXS curve showing the intensity as a function of momentum transfer *s*. The experimental data curve is given as black dots with grey error bars, and the red line is for the GASBOR *ab initio* model fit (*χ*^2^ of 1.14). (ii) The distance distribution, shown as the *p*(*r*) function, provided a maximum particle diameter (*D*_max_) of 17.0 nm. (iii) The Guinier plot revealed a Guinier region (red line) in the range of *sR_g_ <* 1.3, leading to a *R_g_* of 5.35 nm. (iv) The Kratky plot agrees with a compact shape for the mGBP7 dimer with certain flexibility.

##### SAXS

With SAXS we also aimed to reveal the mGBP7 dimer organization. We used a protein concentration range of 1.66–6.85 mg/ml and merged the low *s* data from the 1.66 mg/ml concentration with the high *s* data from the 6.85 mg/ml concentration. The SAXS data are summarized in Table S3. We calculated an *ab initio* dimer model with GASBOR,^62^ using a *P* 2 symmetry, yielding the dimer model shown in Fig. 5C (denoted as model 4 here) which has a tail–tail interface between the E domains of the two mGBP7 chains. We calculated the theoretical intensity of the mGBP7 dimer model with CRYSOL^64^ and compared it with the experimental scattering data, resulting in a *χ*^2^ of 1.89 (Fig. 5D(i)). With the Guinier approximation,^65^ we determined the radius of gyration (*R*_g_) as 5.35 nm and used the distance distribution, shown as the *p*(*r*) function, to specify the maximum particle dimension (*D*_max_) as 17 nm (Fig. 5D(ii) and (iii)). The dimensionless Kratky plot showed a compact elongated shape for the mGBP7 dimer (Fig. 5D(iv)).

Since the *χ*^2^ value for model 4 is somewhat large, we created a further dimer model for mGBP7 (denoted as model 5) using SASREF,^63^ which performs quaternary structure modeling of a complex formed by subunits with known atomic structure against the SAXS data set. The resulting model has larger *R*_g_ and *D*_max_ values compared to model 4, yet the *χ*^2^ is closer to 1 with a value of 1.27 (Table S4). As model 4, this dimer has a tail–tail interface, yet the dimerization is predicted to occur via the M domains (Fig. 5C, bottom). We further tested how good the three ClusPro models (models 1 to 3) are in agreement with the SAXS data. To this end, we docked them into the calculated molecular envelope. Table S4 lists the resulting *χ*^2^, *R*_g_ and *D*_max_ values. A *χ*^2^ close to 1 is desirable, yet also large deviations from the experimentally determined values of *R*_g_ = 5.35 nm and *D*_max_ = 17 nm render a dimer model as disagreeing with the SAXS data. Indeed, more emphasis is given on the agreement for the physical dimensions *R*_g_ and *D*_max_ than on *χ*^2^. Based on these considerations, none of the three mGBP7 models created by ClusPro fully complies with the SAXS data; only model 3 might still be a suitable model. The least fitting is model 1, as its overall length is clearly larger than the SAXS envelope. A head–head interface via the G domains as predicted by AlphaFold (Fig. S7) is for the same reason also unlikely as its *D*_max_ value is in conflict with the calculated *p*(*r*) function.

#### 2.3.4 Structural details of the mGBP7 dimers

In contrast to hGBP1 and mGBP2, mGBP7 shows a strong tendency to dimerize via its stalks in an antiparallel arrangement (Fig. 4). This is seen for models 2 and 3 created by ClusPro, and also applies to the next five best docking predictions not further discussed here, and models 4 and 5 resulting from fitting to the SAXS data. The crossed-stalks motif present in model 2 and to some extent also in model 5 is reminiscent of the dimerization pattern observed for other members of the dynamin superfamily.^66, 67^ Only model 1, the most probable docking prediction, dimerizes via the G domains and some contacts involving *α*13 (Fig. 4). However, this dimerization mode is different from the G domain dimers observed for hGBP1 and mGBP2. This structure is further interesting since it fulfills 3 of the 4 constraints imposed by XL-MS. Regarding the fourth constraint it should be noted that it clashes with the other three constraints, as revealed by a DisVis analysis using the HADDOCK webserver,^68^ indicating that no dimer model would be able to fulfill all four XL-MS constraints at the same time. It is possible that this constraint is due to an artifact, a transient conformational change, or that more than one dimer shape can be assumed. With regard to the dimer interface, model 3 buries the largest protein surface with 3,414 Å^2^, followed by model 1 (2,678 Å^2^) and model 2 (2,581 Å^2^). The smallest contact areas are observed for model 4 (2,224 Å^2^) and model 5 (1,560 Å^2^), which further decreased during the MD simulations. This indicates instability of these models. Moreover, model 4 had to be slightly adjusted to remove atom clashes before the simulation could even be started. Model 2, on the other hand, notably increased its dimer contact area during the simulation, suggesting stability and optimization of the dimer structure. Each mGBP7 dimer model is now discussed in more detail.

##### Model 1

The flexibilities of the M/E domains of mGBP7 in the dimer model 1 differ from the structural fluctuations (measured by the RMSF) of the mGBP7 monomer and are more similar to those of the mGBP2 dimer (Fig. S8). In particular *α*10 and *α*12 are more flexible than in the mGBP7 monomer. The guanine cap is stabilized, involving stronger intramolecular salt bridges between R57 and E254/D255 than in either the apo- or GTP-bound mGBP7 monomer (Fig. 3). Within both chains, *α*4‘ and *α*12 remain connected via a network of four redundant salt bridges of which in either chain two remain highly occupied (Fig. 3). However, the salt bridge R225–E562 completely disappeared. The first principal component of dimer model 1 is a butterfly motion, wherein the hinges move in a correlated fashion to form a more curved dimer, while the CT tail unfurls from its pocket. The second principal component combines the butterfly motion (yet in a reduced from) with a slight screwing motions of the M/E domains. The directions of the hinge motions in both chains are the same as in the hGBP1 and mGBP2 monomers and dimers, with +Δ*y/* + Δ*z* as preferred direction and *−*Δ*y*/+Δ*z* being possible too. The maximum distance of *d*_480_ is 25–31 Å and on average it is *≈*10 Å, which is comparable to the motions in the mGBP7 monomer (Fig. S6). The strongest intermolecular interactions in the mGBP7 dimer model 1 are between the L1 and *α*4’ (E161–K213’, S211–D164’), between *α*4’ and the E domain (G209-K567’, R222–R566’), and between the two E domains (R566–E562’) with interaction energies in the range of *−*80 to *−*40 kJ/mol (Fig. S9). These are partly the same residues as those involved in intramolecular salt bridges, showing the competition between intra- and intermolecular salt bridges, and explaining that the intramolecular salt bridge R225–E562 completely disappeared while those involving R566 or K567 were absent in one of either chain of the dimer.

##### Model 2

Interestingly, the protein motions within model 2 are very similar to those in model 1, even though the dimerization interface is completely different. The M/E domains in models 1 and 2 are similarly flexible. The intramolecular salt bridges in the guanine cap strengthened in both dimers compared to the monomer, while the number of salt bridges formed between *α*4‘ and *α*12 is smaller, but the overall contact remains intact. As in model 1, the R225–E562 salt bridge is not formed in either chain of model 2 and E222–R566 is also not present in chain 1. This is also the chain displaying a higher tip flexibility, and we can thus conclude that the E222–R566 salt bridge strength is anticorrelated with the flexibility of the M/E domain. The butterfly motion resulting from the synchronous hinge motions of both chains is also present in model 2. Nonetheless, the first principal motion is decidedly asymmetric, with the stalks twisting around each other, which results in a contact between one G domain to the stalk tip of the other chain. This can be correlated with the observation of dissolved salt bridges between *α*4‘ and *α*12 and a higher stalk tip flexibility. The second principal motion is similar, albeit symmetric, and here the motion originates not from the stalk-middle but from the base of the G domain. In analogy to DSP nomenclature, this could be termed hinge 2, which is not strongly visible in the GBP monomers. The directions of the hinge 1 motion in model 2 are the same as in model 1 (Fig. S6). The strongest intermolecular interactions are found for the two electrostatic interactions E389–K508’ and E399–R396’ reaching about *−*110 kJ/mol, with supporting weaker interactions around residues 510 and 610.

##### Model 3

In model 3, the mGBP7 proteins are less mobile compared with the other dimer models and also the protein monomer (Fig. S8). Only the CT tail remains flexible. The occupancy of the intramolecular salt bridge in the guanine cap is similar in the mGBP7 monomer and this dimer model, and thus smaller than in the other dimers. Moreover, also three of the four salt bridges tackering *α*4‘ and *α*12 together have an occupation below 50%, indicating an increased tendency of the E domain to detach from the G domain. However, the fourth salt bridge, R225–E562, remains present to a similar degree as in the monomer, whereas in models 1 and 2 this particular salt bridge was absent (Fig. 3B). The first principal motion of model 3 results in a saddle-like structure where the rather planar shape of the dimer becomes more curved due to both stalk tips moving into the same direction. The tip of one of the M/E domains even detaches from the G domain of the other chain, enabling a larger range of motions for the E domains that together almost form a cross shape. The main parts of the helices of the M domains, on the other hand, remain parallel with respect to each other, meaning that the *α*12 helices become slightly angled against the rest of the protein. The motion is accompanied by the CT tails curling up against the G domains. The second principal motion is very similar to the saddle-formation motion, yet with only one of the CT tails curling up, while the other one stays compact. The tips of the M/E domains move mostly in +Δ*y/ −* Δ*z* and *−*Δ*y/* + Δ*z* direction, and the range of the motions is larger than in the monomer (Figs. 4 and S6). The maximum amplitude of that motion is 25–38 Å, with the average values being between 10 and 20 Å, which is comparable to the motions in models 1 and 2. However, all three mGBP7 dimer models discussed so far move generally less than the mGBP2 dimer, which replicates the findings for the mGBP7 and mGBP2 monomer. The intermolecular interactions that contribute the most to the dimer stability are R33–D411’ and K26–E419’, reaching *−*120 kJ/mol and *−*60 kJ/mol, respectively, which are supported by weaker interactions between the guanine cap and the stalk tip (Fig S9). Moreover, the N-terminal E8 is also of relevance as it interacts with several residues of *α*10 of the other chain, summing up to an additional *−*140 kJ/mol of interaction energy.

##### Model 4

Model 4 is generally very flexible, which is caused by the overall instability of this dimer model, involving the loss of one of the two main interprotein contact interfaces (Fig. 4). In terms of intraprotein flexibilities, the SW2, G4 motif, and guanine cap are more mobile than in models 1 to 3, while the *α*10/12 region is only slightly more flexible than in models 1 and 2. The guanine cap is of similar flexibility as in the mGBP7 monomer, which is also reflected in a comparable occupancy of the intramolecular salt bridge within that region. The considerable dissolution of the intramolecular salt bridges connecting *α*4’ and *α*12/13 in chain 2 of model 4 (Fig. 3B) could be at fault for the increased mobility of the dimer, as *α*13 provides a large part in the contact area between both chains. As in model 3, the first principal component can be described as a saddle formation due to a bending of both chains. The second main motion is a sliding of the stalks against each other along the M/E domain axis, which likely results from the accumulation of negatively charged amino acid residues at the dimer interface and the involvement of the highly flexible CT tails therein. Model 4 nominally has a greater range of motions compared to the other dimers, but due to the disruption of the dimer, this is less meaningful. The strongest interprotein contacts are K588–E501’ and E501–R636’ with energies of about *−*50 kJ/mol, which is weaker than for any of the primary interaction pairs in the other dimer models. Moreover, the first principal motion even results in an opening of the contact site around E501-K637’. The conclusion therefore is that model 4 is not a stable dimer.

##### Model 5

The final mGBP7 dimer model displays a crosswise interaction via the top parts of the two M/E domains, which prevents contact formation between the stalk tip of one chain and the G domain of the other chain as in models 2 and 3. However, this interaction interface is not particularly stable as within the 100 ns MD simulation, it is broken and replaced by an asymmetrical stalk tip–G domain interaction. Reasons for that asymmetry are the higher flexibility observed for one of the CT tails and a generally great accumulation of negative charges at the original dimer interface. The motion leading to the change in interaction interface is contained within the first principal component, while the second principal component is an asymmetric motion caused by one chain swinging out and rotating against the other. The individual chains display the same flexibility pattern observed before. The range of the hinge motion is comparable to that in model 3, but as for model 4 this is not of relevance due to instability of model 5. For the latter reason, the overall flexibility of this dimer model is as high as for model 4. The intramolecular salt bridge in the guanine cap is not particularly strong (20–30% population on average), while the *α*4’–*α*12/13 lock is still engaged (50–76% on average). The only strong intermolecular contact that is left from the original interface is E336–R418’, contributing *−*100 kJ/mol.

## 3 Discussion and Conclusion

We compared the sequence and biochemical properties of three GBPs, the human GBP1 and the murine GBP2 and GBP7. The main conclusion from that juxtaposition is that mGBP2 is more similar to hGBP1 than mGBP7 is to mGBP2. Both hGBP1 and mGBP2 are post-translationally modified by isoprenylation where instead mGBP7 exhibits 49 additional C-terminal residues for membrane binding. Moreover, the sequence of mGBP7 differs markedly at various places from those of hGBP1 and mGBP2. The different distribution of charged residues gives rise to different protein dynamics, as we revealed by HREMD simulations of the monomeric proteins. We generated dimer models with ClusPro for the three GBPs, where hGBP1 and mGBP2 involve the G–G domain dimer, also called head-to-head dimer in the following, that was already solved as crystal structure for the G domain of hGBP1.^3^ For mGBP7, we obtained three dimer models, of which model 1 agrees with our XL-MS data. We generated two further mGBP7 dimer models by reconstruction from SAXS data.

### 3.1 General comparison of the dimer models

In order to understand the dimer dynamics, we first simulated the monomers of the three GBPs in solution serving as a reference. For mGBP2 and mGBP7 both the apo- and GTP-bound state were considered, while the hGBP1 monomer was only simulated as apo-protein. In the apo-state of the GBP monomers, the different motifs and loops of the G domains are very flexible, one reason being that R48, which serves as arginine finger, has no stable contact without GTP. The simulations further revealed that all three GBPs feature the characteristic hinge motion involving the M/E domain that we had originally uncovered for hGBP1.^22^ Among them, hGBP1 exhibits the largest hinge motion and mGBP7 the smallest one. A probable reason for this is that the CT tail has a stabilizing effect on the M/E domain of mGBP7. Another difference is that the G domain of mGBP7 does not stabilize so much upon GTP binding as in mGBP2, especially the guanine cap does not adopt a structured conformation and remains flexible as in the apo-state. There are differences in the sequence of the mGBP7 guanine cap compared to hGBP1 and mGBP2. In particular, this motif is negatively charged in mGBP7, which would explain that binding of the negatively charged GTP has no stabilizing effect on it. This in turn further unfolds why none of the three techniques used for determining the mGBP7 dimer structures, i.e., XL-MS, SAXS, and protein–protein docking, predicted the dimerization to occur via the two guanine caps. It is either too dynamic to allow a stable mGBP7 dimer to be formed and/or the amino acid composition does not allow strong enough interactions between the two guanine caps to evolve. In hGBP1 and mGBP2, the stabilization of the guanine cap via GTP binding seems to be a prerequisite for dimer formation, either to reduce the entropy penalty for the dimerization or to have a well defined interaction surface. Since the guanine cap seems not to be involved in mGBP7 dimerization, this further explains why this protein can be a transient dimer in the cytosol without GTP.^24^

Five alternative mGBP7 dimer models where predicted instead, where the protein–protein interface is either between the G domain and *α*12/13 (model 1), between the M/E domains (models 2 and 5), the two M domains (model 3), or the two E domains (model 4). The two mGBP7 models derived from fitting of the SAXS data, models 4 and 5, turned out not to be stable. Model 5 adopted an asymmetric arrangement very quickly. Model 4 might become more stable in the presence of a lipid membrane where binding to the membrane would remove the flexible and this interaction-disturbing CT tail from the interface area.^24^ However, this is a speculation for the time being. Based on the current results, our conclusion is that the mGBP7 dimer structures created by protein–protein docking with ClusPro are more sound. Common observations for these three dimer models are that the intramolecular salt bridges locking *α*4‘ and *α*12/13 never completely dissolved, due to a redundancy of four salt bridges of which at least one is always present with *>*50% occupancy. The G domain dimers (hGBP1, mGBP2, and model 1 for mGBP7) show an increase in the stability of the intramolecular salt bridge within the guanine cap, which coincides with a structural stabilization of the guanine cap, while this salt bridge is not stabilized in the stalk dimers of mGBP7 (models 2 and 3). It should be noted though that the mGBP2 monomer already has a strong salt bridge in the guanine cap. A special observation for the mGBP2 dimer is that in one of the chains all salt bridges between *α*4’ and *α*12 completely dissolved, which on longer time scales could give rise to a folding out of the E domain, as was already observed for hGBP1 multimers.^31, 45, 69^ All dimers are stabilized by intermolecular salt bridges, with one or two of them being of particular strength with interaction energies below *−*100 kJ/mol, which are supported by several surrounding weaker interactions. The biological relevance of the different findings are discussed in the following sections.

### 3.2 Linking structural and mutation data to the current dimer models

#### Dimer interface size

According to Bahadur et al., a biologically relevant dimer interface should comprise at least 1,000 Å^2^.^70^ All six dimer models studied here fulfill this criterion. Negi et al.^71^ had analyzed more than 70 protein complexes with known 3D structures and revealed that dimer contacts are often mediated by „hot spot“ residues, meaning that only very few amino acids significantly contribute towards the binding energy. This is also reflected in our GBP dimer models. The interaction area is largest for the hGBP1 G domain dimer, indicating that this might be the „strong initial interaction“ around which the polymer lattice on a lipid membrane is built.^28^ Previous dimers reported for DSPs (some of which may only be due to crystal-packing) include the hGBP1 dimer by Prakash et al.,^41^ burying 2,890 Å^2^ of surface area for head-to-tail dimerization via G domain–*α*10/12 interactions, and 2,140 ^2^ for the head-to-head arrangement. In 2006, Ghosh et al.^3^ described a hGBP1 dimer that has an even larger interface of 3,900 Å^2^ and includes residues that are conserved in hGBP isoforms and the G domain motifs. The G domain dimer of OPA1, on the other hand, buries only a surface area of 1,573 Å^2^;^10^ however, as the authors did not provide the formula how the interface area was determined, it might be larger by a factor of two. The interfaces buried by our dimers is within or even beyond that range, with values between 5,314 Å^2^ (hGBP1) and 1,560 Å^2^ (mGBP7, model 5). These interfaces thus fulfill the necessary size requirement for a stable dimer. The largest interface sizes were observed for the G domain dimers, which were also identified as the most stable dimers, further confirming the existence of this GBP dimerization mode.

#### Salt bridges at the dimer interface

According to Praefcke et al.,^47^ the residues 48–52 of hGBP1 are necessary for dimerization, especially R48 for building tetramers. Further residues involved, yet to a lesser degree, are 72, 76, 99, 103 106, 109 and 112.^47^ We now understand the relevance of these residues for dimerization due their importance in GTP binding and subsequent stabilization of the G domain, in particular the guanine cap. Moreover, our simulations of mGBP2 also revealed direct engagement in dimerization for two of the residues in that region, namely for R48, which has a small, but noticeable dimerization energy contribution, and E102, which shows a prominent interaction to a number of residues. In a study of hGBP1, where the production of GMP was used as an indirect measure of dimerization, it was demonstrated that the *α*6 helix plays a critical role in dimerization, especially the residues 289–308, as well as residues 103–108.^72^ While the former region was not close enough to the dimer interface in any of the dimers studied here, we can confirm that the latter region has a high energetic contribution to dimer formation of both hGBP1 and mGBP2.

Vöpel et al.^32^ observed that two positively charged residues of hGBP1, R227 and K228, located on *α*4’, as well as four glutamate residues, 556, 563, 568, and 575 located in *α*12/13, form intraprotein salt bridges and that loosening their interaction facilitates dimerization and tetramerization. This can either mean that (i) *α*4’ and/or *α*12/13 are part of the interface causing the intraportein salt bridges to break, (ii) that they need to be broken in order to reveal the interface, or (iii) that dimerization could cause these salt bridges to break via an allosteric effect if *α*4’ and *α*12/13 should not be directly involved in the dimerization. The effect (i) was observed in two of our mGBP7 dimer models, where these salt bridges loosened and either both *α*4’ and *α*12 (model 1) or only *α*12 (model 4) contributes to dimer formation. In another study it was found that deleting the hGBP1 region after residue 481 abolishes tetramerization,^32^ which is broadly reflected in the mGBP7 dimer models 1, 2, and 4, where the region 481–638 is involved in the dimer interface. In 2012, Wehner et al.^54^ suggested that residues 105, 186, 245, and 259 have only minor effects on the dimerization of hGBP1, while residues 240 and 244 (i.e., the guanine cap) where identified to be essential for it. Our results for the head-to-head dimer of hGBP1 revealed R48, E105, K209, D237, D239, R242, K243, R245, K247, E251, K252, E256, E259, and E261 as main players in the dimerization, underlining the importance of the guanine cap. These residues have further in common that they are all charged, and under that aspect, R243 and K244 appear interchangeable. It is interesting to note that all of these residues are conserved between hGBP1 and mGBP2, and often also mGBP7. Although ClusPro did not propose a dimerization of mGBP7 that would correspond to the G domain dimer of hGBP1, we do not want to completely rule out its existence, also considering that AlphaFold-Multimer predicted this as the only mGBP7 dimer model.

In general, we can confirm the importance of residues in the guanine cap (230–245) for dimerization and have indications for the relevance of *α*13 (560–590) for tetramerization. The salt bridge 62–255 that enhances dimerization shows a slightly increased formation in the dimer, and the salt bridge 227–558, which supposedly inhibits tetramerization, shows a slightly decreased probability, especially in dimer models 2 and 4 of mGBP7. Energy analysis of interfacing residues suggests that the intramolecular salt bridges might be replaced by intermolecular interactions.

### 3.3 Biological relevance of different dimer structures

In order to understand why different GBP dimer structures may be adopted, it is instructive to recall their conformational dynamics. Upon dimerization, the hinge motions of the two proteins forming the dimer give rise to an overall correlated motion. In the cases of the mGBP2 dimer and model 1 of mGBP7, the two chains do not show a greater individual range of motions than their monomeric counterparts, but the extended length of the two proteins moving in a concerted fashion could introduce more stress on a membrane than a single GBP protein is able to do, comparable to the observations made for BAR (Bin/Amphiphysin/Rvs) domain proteins.^73–75^ In contrast to that, the motions of dimer model 2 of mGBP7 do not lead to an increased overall curvature, so that a possible mechanism of introducing membrane stress/shearing force would require both the CT tails and the stalk tips to be inserted into the membrane. The insertion of a single (angled or short) helix can result in a hydrophobic mismatch that facilitates membrane fission,^76^ comparable to a wedge driven into the lipid bilayer. The dimer models 3 and 4 of mGBP7 involve a more flexible stalk tip than in the monomeric mGBP7 form, but the protein–protein interaction along the length of the M/E domains constricts the overall dynamic effect.

While these three notable structures seem initially to be at odds with each other, several studies of other DSPs are in support of the existence of different DSP multimer arrangements based on up to four intermolecular interfaces.^75, 77^ In the case of OPA1, dimerization can be both nucleotide-dependent and -independent, involve multiple interfaces apart from the G domain, and lead to higher order oligomers.^10^ It is possible that the structures spatially complement each other to form a lattice, a multimer consisting of several different dimers. On the other hand, the dimers may also temporally complement each other during the GTPase cycle, with rearrangements triggered by nucleotide binding or hydrolysis. Moreover, some dimer structures may not be functional and only be present during the aggregated/storage phase.

If we translate the structural information that is available for DSP multimers to the GBPs studied here, we conclude that two of the five mGBP7 dimer models (models 1 and 5) can be combined to form a long string of half-moon shapes, comparable to the BAR domain proteins (Fig. 6). The primary, most stable interaction would happen via the G domains, with the underside of the E domains forming the second interface. The alternating rise and fall of the proteins could induce local curvature of the membrane, especially when considering the hinge motion of the M/E domains. Adding a third model (model 3 or 4), the dimers can be integrated into a helical-ring-model, reminiscent of other dynamins. While the first turn of the helix consists of G and E domain interactions, the remaining M domain would form the inter-rung contacts. A conformational change of the M/E domain would be needed to „unlock“ the fourth interface, the crossed-stalks dimer. This further requires that the G domain interface is flexible enough that a tilting of the spherical G domains against each other is still stable. The docking results indicate that such non–standard G domain interfaces are possible. Such a ratchet-like tightening of the dynamin collar has been previously suggested.^2^

**Figure 6:**
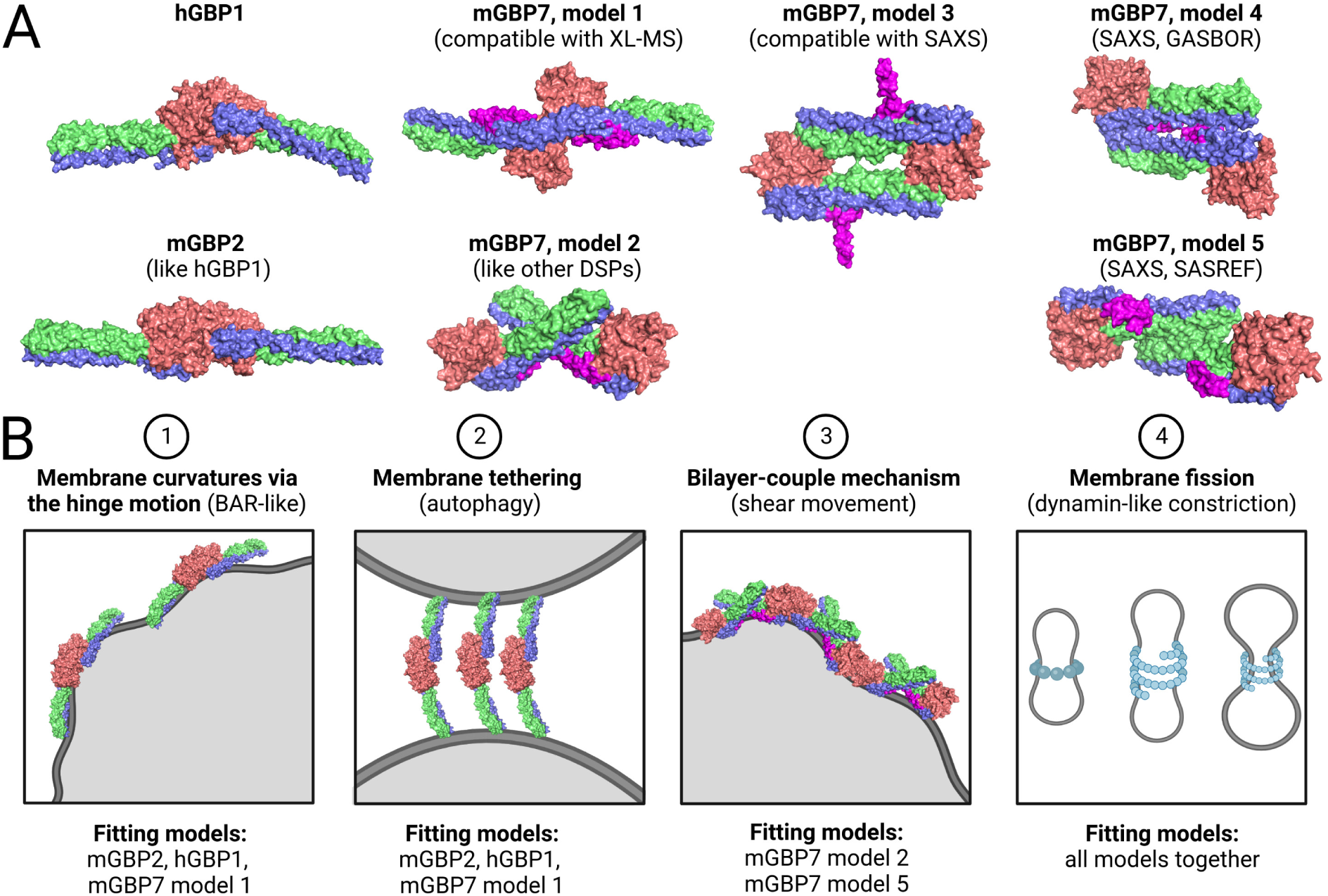
Summary of dimer structures: **(A)** Seven dimer models of hGBP1, mGBP2, and mGBP7 were studied in this work. **(B)** Different dimerization modes suggest various mechanisms of action. A curving of the dimer could induce membrane curvature by the scaffold mechanism, similar to BAR domain proteins or the COP (cytoplasmic coat proteins for vesicle transport) machinery. The G domain dimer of hGBP1, mGBP2, or model 1 of mGBP7 could serve for tethering the parasitophorous vacuole membrane to the autophagosome. Insertion of tilted helices, which would be possible with mGBP7 dimer models 2 and 5, would give rise to the so-called local spontaneous curvature mechanism or bilayer-couple mechanism^73^ and ruffle the membrane or generate shearing forces. Finally, by combining several of these dimerization modes, even a dynamin-like constriction would be imaginable. This figure was created with BioRender.

Furthermore, with more than one interface, the dissolution of less stable interactions could be prevented by a lattice of surrounding interactions, and additional binding of the CT tails to the membrane. The small sequence variations between hGBP1, mGBP2 and mGBP7 seem indeed to be sufficient to tip the scale in favor of other dimer structures. It might thus be possible that some of the isoforms act as chain-starters (with maybe only one stable interface), some as continuous elongators, and some as chain-breakers (with no further interfaces), a scenario in which heterooligomers of seemingly redundant proteins would be necessary. For bacterial DSPs, this has already been proposed.^33^ We further know that mGBP2 and mGBP7 co-localize at the parasitophorous vacuole membrane and that mGBP2 attaches first, followed by mGBP7, and that mGBP7 deletion is more lethal than mGBP2 deletion.^17^ Perhaps mGBP7’s various polymerization options allow a larger carpet of GBPs to form, which can then more effectively remodel membranes, something that should be addressed by future studies.

## 4 Methods and materials

All details about the experimental methods are provided in the Supplementary Information.

### 4.1 MD simulations and docking

#### General aspects of the MD simulations

The structures of hGBP1, mGBP2, mGBP7 were prepared for the simulations as previously described, which includes the parametrization of GTP and the geranylgeranyl group.^22–24^ The hGBP1 dimer was created using MMa-lign (https://zhanglab.ccmb.med.umich.edu/MM-align/^58^). For all MD simulations, the Amber99SB*-ILDNP force field^78–80^ was used for modeling the proteins, the TIP3P model was used for water,^81^ and GROMACS 2016^82, 83^ was employed for running the simulations. The simulation temperature and pressure were 310 K (37 *^◦^*C) and 1 bar, respectively. In Table 2, all MD simulations included in this study are listed.

#### HREMD simulation of the mGBP7 monomer

In our previous study of mGBP7,^24^ we already obtained the structure of mGBP7 by using homology modeling and simulated it as mGBP7_GTP_ for 100 ns. The most populated cluster structure of that simulation was used here as starting point for the HREMD simulations. To identify the role of GTP and the CT tail, two further systems, mGBP7_GTP_ and mGBP7_holo_ were created, using an identical approach as described on our work of mGBP2.^23^ The protein was always placed in a rectangle simulation box of 10 nm *×* 10 nm *×* 18.0 nm dimensions, ∼56,000 water molecules were added, as well as 10 (apo), 13 (GTP), or 13 Na^+^ (holo) for the neutralization of the systems, resulting in a total number of ∼178,000 atoms. A similar HREMD protocol as applied in our study of mGBP2 dynamics was used and is described in detail in the Supplementary Information.

#### Protein–protein docking

To obtain GBP dimer structures, we performed protein–protein docking and employed ClusPro (https://cluspro.bu.edu)^56^ for this purpose, taking favorable assessments of this program into account.^85, 86^ The multimer docking mode of ClusPro was used, both in combination with SAXS data and/or crosslinking data, and without additional restraints. The inclusion of additional restraints did not result in new structures, only a reordering of the results happened. From the top 10 results of the different setups, the most promising candidates were selected for MD simulations.

#### MD simulations of the GBP dimers

The dimers were simulated in boxes with sizes, depending on the structure, varying from 27.0 nm *×* 15.5 nm *×* 13.5 nm to 18.5 nm *×* 12.5 nm *×* 13.5 nm, with water added for solvation and NaCl added for neutralization. A two-step equilibration was performed as described for the HREMD simulation, yet without an additional *NpT* equilibration. The hGBP1 and mGBP2 dimers were simulated with GTP being bound, using a 2 fs time step, while the GTP-free mGBP7 dimers were simulated using a 4 fs time step. The reason for simulating mGBP7 in its apo-form is two-fold: firstly, it was shown by Legewie et al.^24^ that mGBP7 can form dimers in the absence of GTP; and secondly, without GTP, virtual sites can be applied enabling a larger time step. In all simulations, distance restraints of 1,000 kJ mol*−*1 nm*^−^*^2^ were applied between the *β*-sheets of the G domains, in order to inhibit overall rotation, allowing us to keep the box size smaller. The Parrinello-Rahman barostat and Nosé-Hoover thermostat were used. Each dimer structure was simulated for 100 ns, where frames were saved every 20 ps and analyzed. In order to assess the dimer–dimer interactions, another 100 ns MD trajectory was produced per dimer, where additional energy groups had been added to the GROMACS index file. For this, the first 100 ns trajectory was analyzed, and all residues within 7.5 Å of the other chain were included. This allowed us to identify the most strongly interacting residues in the second trajectory, which were energetically analyzed in detail.

### 4.2 Analysis of the MD simulations

To generate figures of the 3D protein structures, we applied PyMol.^87^ If not stated otherwise, the analyses were performed using GROMACS 2016. For the analysis of the HREMD simulations, only the target replica was used.

#### Sequence alignment

The alignment of the sequences of hGBP1, mGBP2, and mGBP7 was done with T-Coffee^51^ using the default settings as available at the European Bioinformatics Institute (EMBL-EBI) webserver^52^ (https://www.ebi.ac.uk/services). The resulting sequence alignment was visualized using Jalview 2^88^ where we colored the residues according to their residue type.

#### Root mean square fluctuations (RMSF)

For quantifying the flexibility of the GBPs, we calculated the RMSF of the C*_α_* atoms around their average positions for each residue. A residue with a value over 2 Å is considered as flexible. For visualization, the RMSF values were color-mapped and projected onto the corresponding start structure of the simulations using red colors for flexible regions (RMSF ≳ 2.5 Å) and blue colors for rigid parts (RMSF ≲ 1.5 Å).

#### Electrostatic potential surface (EPS)

The Adaptive Poisson-Boltzmann Solver as available as plugin APBS 2.1 in PyMol was used for the calculation of the electrostatic potential.^89^ To generate the input files (*\*.pqr* and *\*.pot*) needed for this, the web server https://server.poissonboltzmann.org with default settings and with with pH 7 for the *pK*_a_ calculation was employed.^90^ For visualization, the electrostatic potential was color-mapped between *−*5 (red) and +5 kT/e (blue) and projected onto the surface presentation of the proteins.

#### Movements of residue 480

To quantify and characterize the hinge motion of the M/E domain, the motions of residue 480 were monitored relative to the start structure of the simulations. For this, the changes in the Cartesian coordinates (Δ*x*, Δ*y*, Δ*z*) as well as as the absolute distance (*d*_480_) of that residue were determined using our own VMD^91^ script. To do this, we fitted the trajectory frames to the G domain of the start structure. To illustrate the hinge motion, we computed the spatial distribution of residue 480 using the *gmx spatial* tool of GROMACS.

#### Principal component analysis (PCA)

To extract the main motions of the proteins, principal component analyses as implemented in GROMACS were performed. The covariance matrix was created with *gmx covar*, and the first two eigenvectors, also called principal components or principal motions (PC1 and PC2), were determined using *gmx anaeig*. The trajectory was fitted to the C*_α_* atoms, and for the generation of conformations along eigenvectors 1 and 2, the backbone was selected.

#### Salt bridge analysis

To determine the occupancies of selected salt bridges, the distances between the atoms involved were determined. Following atoms of charged residues were used for the calculations: Lys – NZ; Arg – NE, NH1, NH2; Asp – OD1, OD2; Glu – OE1, OE2. A salt bridge was assumed to be present if the distance in question was within 4.5 Å. Salt bridge occupancies were calculated as time-averaged probabilities.

#### Clustering analysis

To characterize the flexibilities of the G domains, we performed clustering analyses for the G motifs and loops using the algorithm of Daura et al.^92^ as implemented in GROMACS. The clustering was applied to all atoms of the desired structural element with a C*_α_*-RMSD cutoff of 2.5 Å to identify cluster membership. Before the clustering, the trajectory was fitted onto the *β*-sheets in the G domain.

#### Protein–protein interface size

The size of the protein-protein interfaces was calculated using the difference in the solvent-accessible surface areas (SASA):

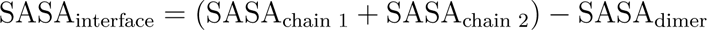

For the SASA calculation, the GROMACS tool *gmx sasa* was used, with a probe radius of 1.4 Å.

#### Analysis of dimer interaction energies

The energy groups of interacting residues as determined in the 100 ns MD simulations of the GBP dimers were added to the GROMACS index files and a second 100 ns MD simulation per dimer was performed. The resulting residue– residue energies were presented as a matrix, and the most negative energies were further analyzed by decomposing them into Coulomb and Lennard-Jones short-ranged interaction energies as calculated by GROMACS.

## 5 Funding and Acknowledgments

This work was funded by the Deutsche Forschungsgemeinschaft (DFG, German Research Foundation)-project number 267205415-CRC 1208 project A07 (J.L., W.S, B.S) as well as and Z01 (K.S.). J.L., W.S and B.S gratefully acknowledge the computing time granted through JARA-HPC (project JICS6A) on the supercomputer JURECA at the Forschungszentrum Juelich.^84^ The Center for Structural studies is funded by the DFG (Grant number 417919780 and INST 208/761-1 FUGG to S.S.). The authors gratefully acknowledge the Gauss Centre for Supercomputing e.V. (www.gauss-centre.eu) for funding this project by providing computing time on the GCS Supercomputer SuperMUC-NG at Leibniz Supercomputing Centre (www.lrz.de). W.S. gratefully acknowledges funding received from Jürgen Manchot Promotionsstipendium. Additional computational infrastructure and support were provided by the Centre for Information and Media Technology at Heinrich Heine University Düsseldorf.

## 6 Data availability

We uploaded the SAXS data to the Small Angle Scattering Biological Data Bank (SAS-BDB),^93, 94^ with the accession code SASDLY9. The mass spectrometry proteomics data have been deposited to the ProteomeXchange Consortium via the PRIDE^95^ partner repository with the dataset identifier PXD026979.

## 7 Author Contribution Statement (draft)

W.S. performed, analyzed and wrote the GBP dimer part. J.L. performed, analyzed and wrote the GBP monomer part. G.P. performed, analyzed and wrote the XL-MS part. J.R. and S.S. performed, analyzed and wrote the SAXS part. D.D. and K.P. provided infrastructure, supervision and protein material. K.S. provided infrastructure and supervision. B.S. conceived the research project and provided infrastructure and extensive supervision.

