## Supplementary Methods, Tables and Figures for "Integrative modeling of guanylate binding protein dimers"

△*Contributed equally to this work.*

### 1 Methods and materials

#### 1.1 HREMD simulation of mGBP7 monomer in solution

The starting structure for all replicas was the same and it was prepared in the following way. The homology model of mGBP7 was solvated and $Na^+$ ions were added for the neutralization of the system. An energy minimization of the solvated system was then performed using the steepest descent algorithm, followed by three equilibration MD simulations. First, equilibration in an $NVT$ ensemble was carried out for 0.1 ns, and then an $NpT$ equilibration for 1 ns position took place. In both of these equilibration steps, the protein atoms were restrained to their positions with a force constant of 10 kJ mol$^{-1}$ Å$^{-2}$ to equilibrate the solvent around the protein and to reach the temperature of 310 K and the pressure of 1 atm. The final equilibration, which was for 20 ns in an $NpT$ ensemble, was without position restraints on most parts of the proteins, apart from the rigid $\beta$-sheets of the G domain. These restraints were kept, also in the HREMD simulation, to avoid overall rotation and translation of the highly prolate protein, which would otherwise require a significantly larger simulation box.[1,2] Thereafter, the HREMD simulations[3] with 30 replicas each were performed for mGBP7$_{\mathrm{apo}}$ and mGBP7$_{\mathrm{GTP}}$. The protein was treated as hot region by modifying its energy function, including the mGBP7–water interactions. To this end, a biasing factor of 310 K/$T$ was applied to each of the 30 replicas, where $T$ is the temperature of the replica in question and which were exponentially distributed between 310 and 450 K. This includes one unbiased replica, called target replica at 310 K. Exchanges between neighbored replicas were attempted every 2 ps, and an average exchange probability of ∼30% was reached. Each replica simulation was 400 ns long, which leads to an accumulated simulation time of 3.2 $\mu$s per HREMD simulation.

To ensure that GTP stayed in its binding pocket in mGBP7$_{\mathrm{GTP}}$, we applied distant restraints between GTP and K51, Y53, and D97 using the pull code of GROMACS. The HREMD simulations were conducted with GROMACS 2016.4 in combination with the PLUMED plugin (version 2.4.1 from `https://github.com/GiovanniBussi/plumed2/tree/`

v2.4).[4] For the temperature and pressure regulation, a velocity rescaling thermostat with canonical sampling[5] and an isotropic Parrinello-Rahman barostat[6] were used. The particle-mesh Ewald (PME) method[7,8] was applied for the calculation of electrostatic interactions in conjunction with periodic boundary conditions. The cutoff value of the Lennard-Jones and short-range electrostatic interaction was 12 Å. We used the leapfrog stochastic dynamics integrator for the integration of equations of motion and the LINCS algorithm[9] to constrain all bond lengths. For the mGBP7$_{\text{apo}}$, we treated certain hydrogen atoms as virtual interaction sites, which permitted an integration time step of 4 fs while maintaining energy conservation.[10] In case of mGBP7$_{\text{GTP}}$, a time step of 2 fs was applied. The coordinates and velocities were saved every 20 ps.

#### 1.2 Protein expression for mGBP7

The expression and purification of the mGBP7 protein based on the protocol of Legewie et. al 2019.[11] For the expression of the mGBP7 protein, competent *E. coli* Rosetta 2 (DE3) pLysS (Novagen) cell were transformed with the pQE-80L vector (Qiagen), containing the n-terminal hexa-histidine tagged mgbp7 gene.[12] We prepared 4 L 2YT (16 g/l tryptone, 10 g/l yeast extract, 5 g/l sodium chloride) media, supplemented with 3.4 $\mu$g/ml chloramphenicol and 10 $\mu$g/ml ampicillin. We inoculate the media with the mGBP7 expression cell with a starting $OD_{600}$ of 0.1 and incubate them at 37 °C and 180 rpm shaking to an $OD_{600}$ of 0.5. The protein expression was started by induction with 150 $\mu$M IPTG and further incubation at 37°C and 180 rpm for 4h. The cells were harvested at 8000g for 30 min and the supernatant was discarded. The resulting cell pellet was resuspended with buffer (50 mM Tris pH 8.0, 300 mM NaCl, 5 mM MgCl, 10 mM Imidazol, 1 mM DTT and 0.25 mM Pefabloc). The cell disruption was done with 2.7 kbar on a Constant Cell Disruption System in 3 cycles followed by a high spin centrifugation step (100000 g, 1h, 4°C). The supernatant was incubated overnight with Ni-NTA-Agarose beads (Qiagen) at 4°C. After the overnight incubation, the Ni-NTA-Agarose beads were washed four times with wash buffer (50 mM

Tris pH 8.0, 150 mM NaCl, 5 mM MgCl, 10 mM Imidazol, 1 mM DTT) and the final elution was done with elution buffer (50 mM Tris pH 8.0, 150 mM NaCl, 5 mM MgCl, 300 mM Imidazol, 1 mM DTT). The elution fraction was concentrated to 5 ml with a 50 kDa cut-off filter and loaded onto a HiLoad® 26/600 Superdex® 200 pg column (Cytiva), preequilibrated with SEC buffer (50 mM Tris pH 8.0, 5 mM MgCl, 2 mM DTT). Elution peak fraction were concentrated and used for further experiments.

#### 1.3 Cross linking mass spectrometry of mGBP7

Here, 6 $\mu$g of mGBP7 was crosslinked in a final volume of 10 $\mu$l using 0.5 mM bis(sulfosuccinimidyl) suberate (BS3) or 0.5 mM disuccinimidyl sulfoxide (DSSO) for 30 minutes at room temperature in 4-(2-hydroxyethyl)-1-piperazineethanesulfonic acid containing aqueous buffer. For control reactions, the crosslinkers were omitted. The reaction was stopped by adding an aqueous solution of 0.5 $\mu$l 1 M tris(hydroxymethyl)aminomethane pH 7.5 for 15 minutes. Samples were separated in 4-12% Bis-Tris polyacrylamidegels. After staining with Coomassie blue, mGBP7 monomer and dimer containing bands were cut-out and processed for mass spectrometric analysis essentially as described.[13] Briefly, the protein was reduced with dithiothreitol, alkylated with iodoacetamide and digested with trypsin. Resulting peptides and crosslinked peptides were resuspended in 0.1% trifluoroacetic acid and analyzed by liquid chromatography coupled mass spectrometry. Peptides were separated for one hour on an Ultimate 3000 Rapid Separation Liquid Chromatography system on a 25 cm length C18 column as described[14] and analyzed by a Fusion Lumos mass spectrometer, online coupled via a nano-electrospray interface.

BS3 crosslinked samples were analyzed as follows: A survey spectra was recorded in the Orbitrap analyser (scan range 400–1800 m/z, resolution 60000, maximum injection time 50 ms, AGC target 100000) and subsequently, 2-10 fold charged precursors were selected (minimum intensity 50000, maximum intensity 1E20, 1.6 m/z isolation window), fragmented with collisional induced dissociation (CID) and independently with higher energy collisional dissociation (HCD). Fragment spectra were recorded in the Orbitrap (resolution 30000, maximum injection time 100 ms, AGC target 50000). The cycle time was set to 2 seconds and already fragmented precursors were excluded from further isolation for the next minute. For DSSO crosslinked samples, survey scans were carried out with following parameters: scan range 350–1600 m/z, resolution 60000, maximum injection time 50 ms, AGC target 400000. Next, 3–8 fold charged precursors were selected (minimum intensity 20000, maximum intensity 1E20, 1.6 m/z isolation window), fragmented by CID (collision energy 25%) and analyzed in the Orbitrap (resolution 30000, maximum injection time 100 ms, AGC target 50000). Subsequently, MS3 scans of two MS2 precursors were carried out in the ion trap for masses matching DSSO induced differences (isolation window 2.5 m/z, MS2 isolation window 2 m/z, scan rate: rapid, maximum injection time 120 ms, AGC target 20000). Finally, an MS2 scan was carried out in the Orbitrap after ETD fragmentation (isolation window 1.6 m/z, resolution 50000, maximum injection time 150 ms, AGC target 200000). The cycle time was set to 4 seconds.

Data analysis of BS3 crosslinked samples was carried out using the mGBP7 amino acid sequence with MeroX (version 2.0.2.4)[15] considering $C_8H_{10}O_2$ as mass shift for the crosslinks between lysine residues. Methionine oxidation was considered as fixed and cysteine carbamidomethylation as variable modification and up to three missed tryptic cleavage sites. Precursor precision was set to 5 ppm and fragment precision to 10 ppm. Crosslinked peptides were reported at a false discovery rate of 1%. For analysis of DSSO crosslinks, the Proteome Discoverer Software (version 2.3.0.523) including XlinkX was used applying tryptic cleavage specificity with a maximum of two missed cleavage sites, carbamidomethylation on cysteines as fixed and methionine oxidation as variable modifications. Spectra associated with potentially crosslinked peptides were filtered (XlinkX detect, +158.004 Da crosslink modification between lysines) and searched by XlinkS (precursor mass tolerance 10 ppm, Orbitrap fragment spectra mass tolerance 20 ppm, ion trap fragment mass tolerance 0.5 Da). Non-crosslinked peptides associated spectra (CID and EThdD) were subjected to a Sequest

HT based searches including hydrolyzed DSSO (+176.014 Da) at lysine residues as additional variable modification. Precursor tolerances were 10 ppm and fragment spectra tolerances 0.02 Da. Identified peptides and crosslinks were accepted at a false discovery rate of 1%. Only crosslinks were reported which were identified in two independent experiments of a sample group.

#### 1.4 SAXS of mGBP7

We performed the small-angle X-ray scattering (SAXS) measurements of mGBP7 on our Xeuss 2.0 Q-Xoom system (Xenocs). This system is equipped with a GENIX 3D CU Ultra Low Divergence x-ray beam delivery system (Xenocs) and a PILATUS 3 R 300K detector (Dectris). The chosen sample to detector distance for this experiment was 0.55 m, results in an achievable q-range of 0.05 – 6.5 $nm^{-1}$. The measurement was performed at 10°C with a protein concentration range of 1.66 – 6.85 mg/ml. The system autosampler injected the mGBP7 samples in the Low Noise Flow Cell (Xenocs). We collect six frames with an exposer time of ten minutes/frame and scaled the data to absolute intensity against water. The radial averaging of the scattering data was done with Foxtrot (v.3.4.9, Soleil/Xenocs). All other used programs for data processing were part of the ATSAS Software package (Version 3.0.3).[16] Primary data reduction (merging of data and background subtraction) was performed with the program PRIMUS.[17] The forward scattering $I(0)$ as well as the radius of gyration ($R_g$) was determined with the Guinier approximation.[18] The program GNOM[19] was used to estimate the maximum particle dimension ($D_{max}$), based on the pair-distribution function $p(r)$. Low resolution ab initio models were calculated with GASBOR[20] with a P2 symmetry. Rigid body modeling of the mGBP7 dimer was done with SASREF.[21] Superimposings of the mGBP7 dimer models were done with the program SUPCOMB.[22] The agreement of the mGBP7 dimer models were checked with the program CRYSOL.[23] The complete data are summarized in Tab. S3.

#### 2 Supplementary information figures

A

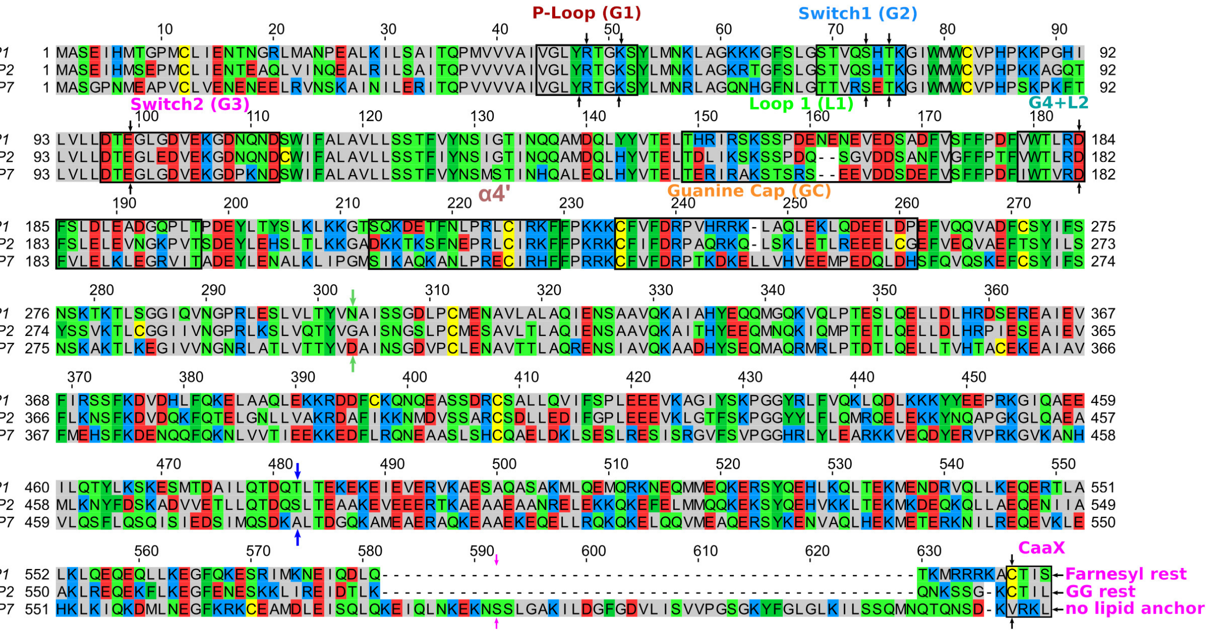


B

| | | aromatic/ hydrophobic | polar | positively charged | negatively charged |
|---|---|---|---|---|---|
| **P-L (G1)** | hGBP1 | 5 | 2 | 2 | - |
| | mGBP2 | 5 | 2 | 2 | - |
| | mGBP7 | 5 | 2 | 2 | - |
| **SW1 (G2)** | hGBP1 | 1 | 5 | 2 | - |
| | mGBP2 | 1 | 5 | 2 | - |
| | mGBP7 | 1 | 4 | 2 | 1 |
| **SW2 (G3)** | hGBP1 | 5 | 4 | 1 | 5 |
| | mGBP2 | 4 | 4 | 1 | 6 |
| | mGBP7 | 6 | 2 | 2 | 5 |
| **L1** | hGBP1 | 6 | 7 | 4 | 7 |
| | mGBP2 | 8 | 8 | 2 | 4 |
| | mGBP7 | 5 | 5 | 4 | 7 |
| **G4+L2** | hGBP1 | 9 | 4 | 1 | 4 |
| | mGBP2 | 9 | 4 | 2 | 3 |
| | mGBP7 | 10 | 2 | 3 | 3 |
| **α4'** | hGBP1 | 7 | 4 | 4 | 2 |
| | mGBP2 | 6 | 3 | 6 | 2 |
| | mGBP7 | 8 | 3 | 5 | 1 |
| **GC** | hGBP1 | 12 | 2 | 6 | 6 |
| | mGBP2 | 12 | 4 | 5 | 5 |
| | mGBP7 | 12 | 2 | 5 | 8 |
| **M domain** | hGBP1 | 80 | 36 | 27 | 29 |
| | mGBP2 | 87 | 40 | 21 | 21 |
| | mGBP7 | 77 | 42 | 29 | 19 |
| **α12** | hGBP1 | 24 | 21 | 16 | 19 |
| | mGBP2 | 28 | 15 | 19 | 21 |
| | mGBP7 | 27 | 18 | 17 | 19 |
| **α13** | hGBP1 | 10 | 8 | 7 | 3 |
| | mGBP2 | 10 | 7 | 6 | 4 |
| | mGBP7 | 10 | 5 | 5 | 5 |
| **CT tail** | mGBP7 | 27 | 13 | 6 | 3 |

**Figure S1: Sequence alignment and characteristics of hGBP1, mGBP2, and mGBP7. (A)** In the sequence alignment, the residues are colored based on their physicochemical properties: apolar, gray; polar, green; aromatic, dark green; negatively charged, red; positively charged, blue; cysteine, yellow. The four conserved GTP-binding site motifs and other important structural elements are highlighted by black boxes and labeled. The key residues for GTP binding and hydrolysis are indicated by black arrows, while the green and blue arrows mark the beginning of the M and the E domain, respectively. The magenta arrows indicate the start of the CT tail in mGBP7. The sequence alignment was calculated with T-coffee[24,25] and the figure created with Jalview 2.[26] **(B)** The count of different residue types in the G motifs of hGBP1, mGBP2, and mGBP7.

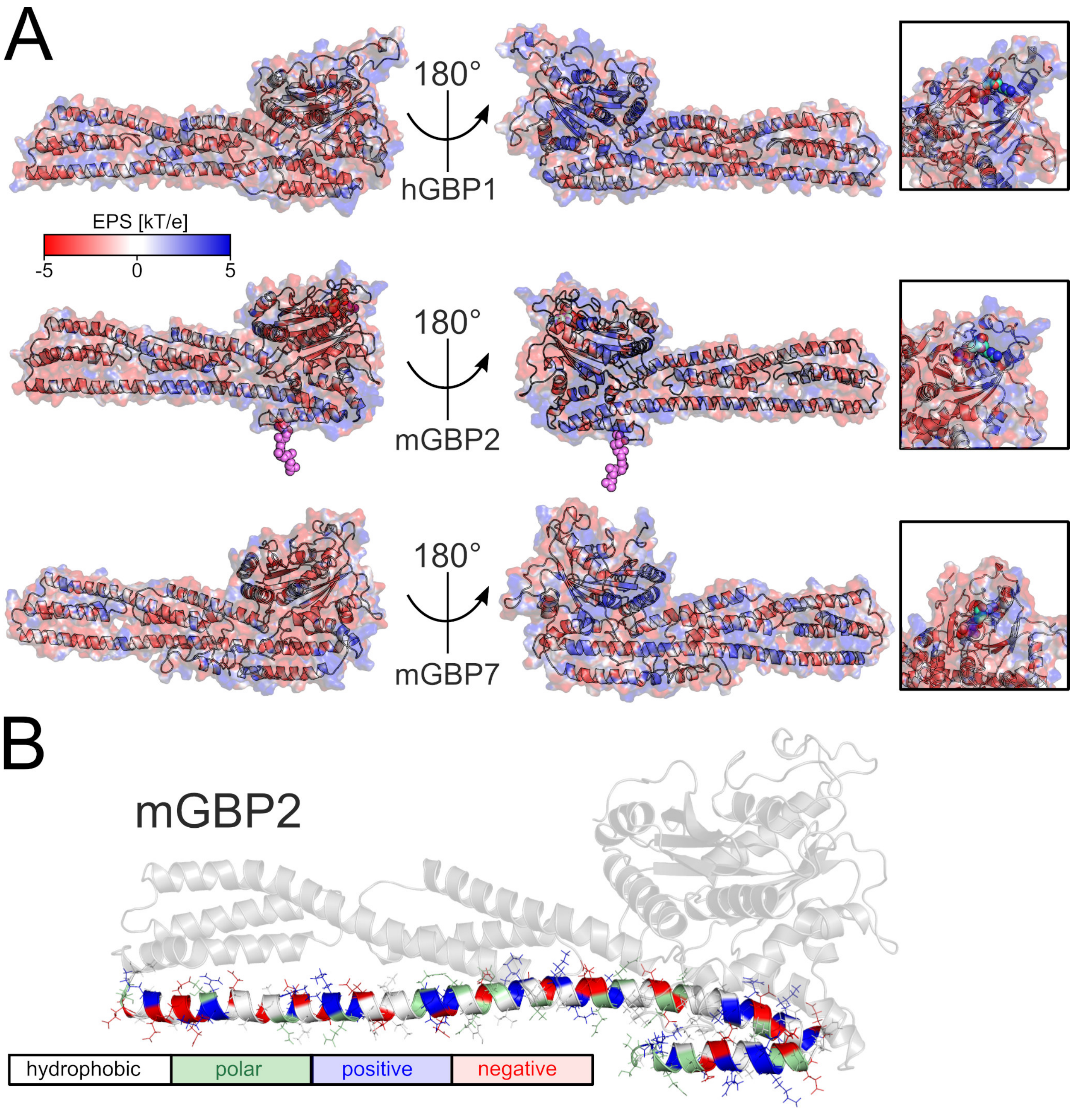


**Figure S2: Electrostatic potential surface (EPS) of hGBP1, mGBP2, and mGBP7 and the distribution of charged residues in the E domain of mGBP2.** **(A)** The EPS was calculated with the APBS webserver and was illustrated with PyMOL and the APBS tool 2.1. The EPS is shown between $-5$ (red) and $+5$ kT/e (blue) for the whole proteins and as zoom for their GTP binding sites. **(B)** The residues of $\alpha 12/13$ of mGBP2 are colored based on their physicochemical properties: apolar and aromatic, white; polar, green; negatively charged, red; positively charged, blue.

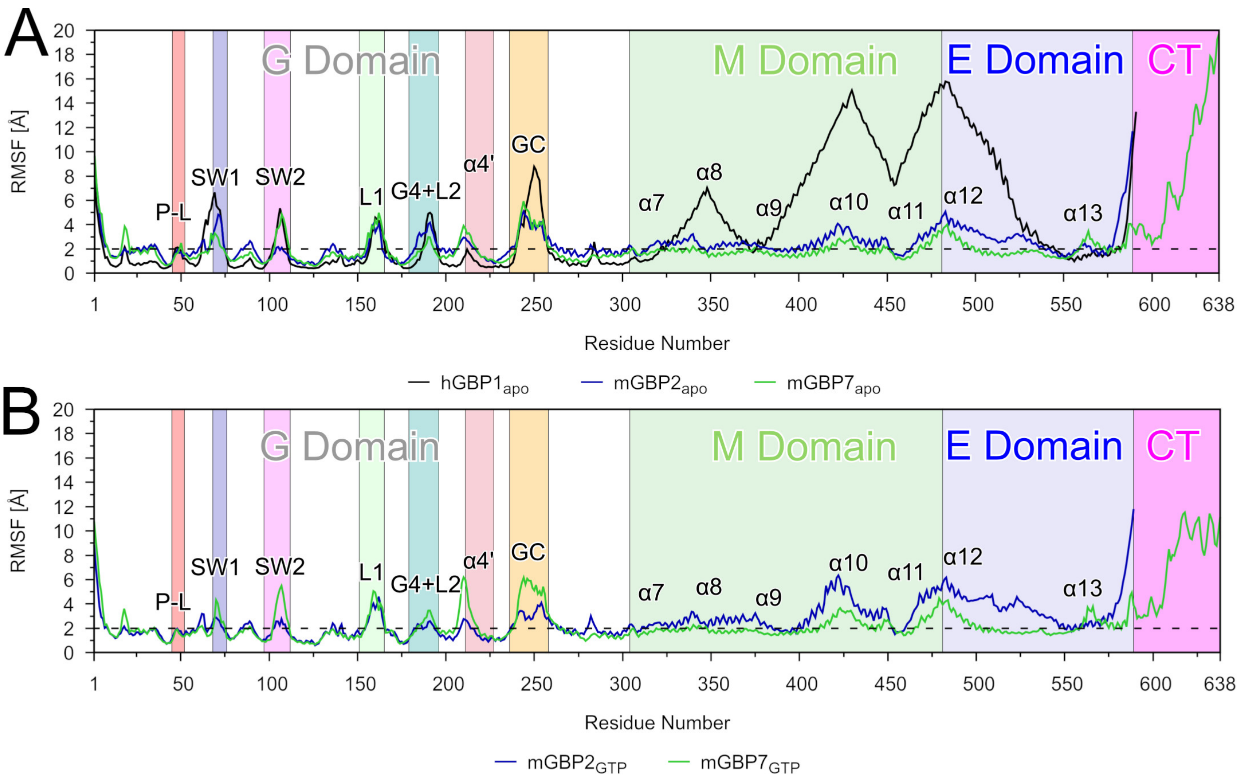


**Figure S3: Fluctuations of the GBP residues during HREMD simulations of the protein monomers.** The fluctuations are quantified by the RMSF of the $C_\alpha$ atoms of hGBP1 (black), mGBP2 (blue), and mGBP7 (green) in **(A)** the apo-state and **(B)** the GTP-bound state. All motifs and loops of the G domains as well as helices in the M/E domain are labeled, and the background of the plots is colored to indicate the different structural parts of the GBPs (where the same colors as in Fig. 1C/D were used). The horizontal dashed line at 2 Å is to identify flexible residues with RMSF values exceeding that value.

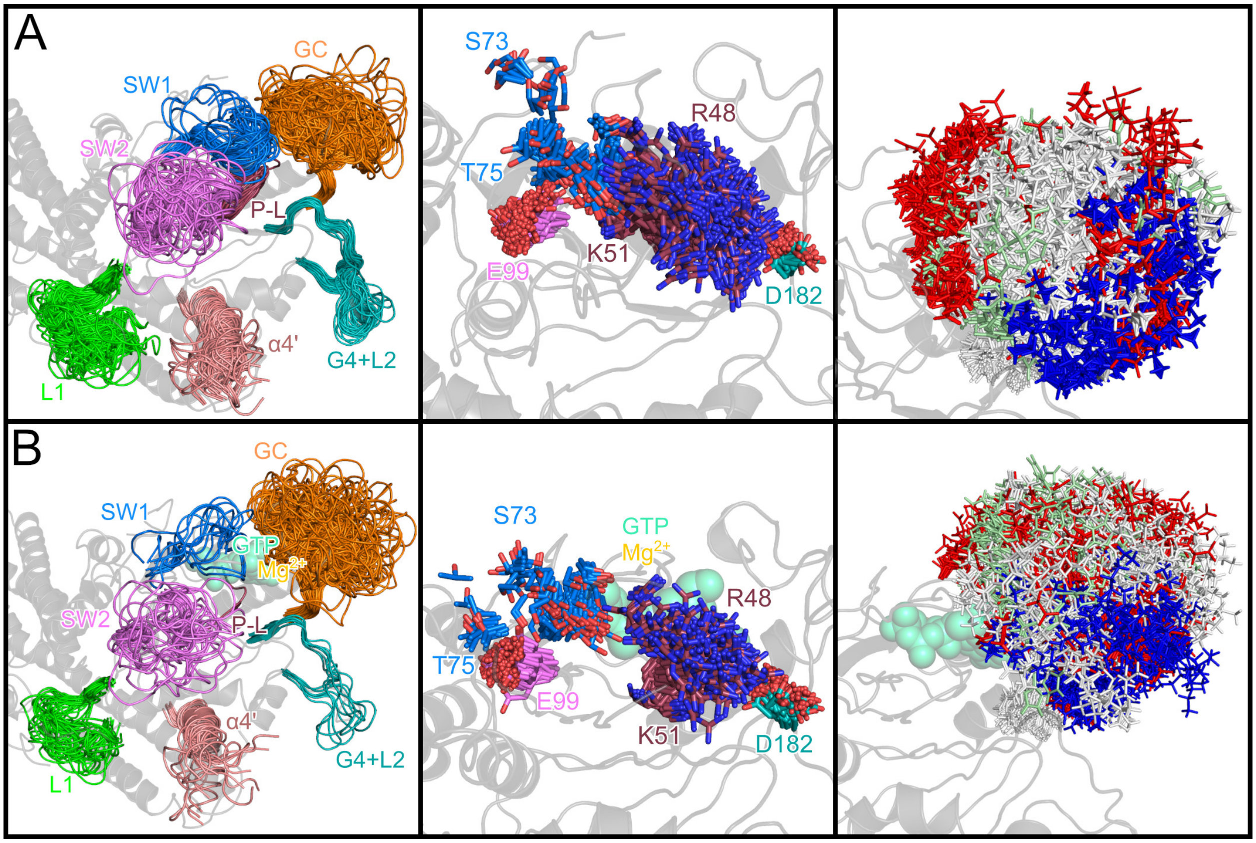


**Figure S4: Conformational clusters of G domain structural elements and selected residues of (A) mGBP7$_{apo}$ and (B) mGBP7$_{GTP}$ monomers.** (Left) Representative structures of the clusters of the G motifs and loops that were determined using a 2.0 Å RMSD cutoff applied to the fitted target replica of the respective HREMD simulation. The colors of the loops are: P-L in red, SW1 in blue, SW2 in magenta, L1 in green, G4+L2 in turquoise, the guanine cap in orange, and $\alpha$4' in salmon. (Middle) The different conformations of key residues for GTP binding and hydrolysis, as sampled during the HREMD simulations, are shown as sticks for the side chains. The oxygen and nitrogen atoms of these side chains are colored in red and blue, respectively, while all other side-chain atoms are shown with the same color as chosen in (A) for the structural element they belong to. (Right) The different conformations of all side chains of the guanine cap are shown as lines and colored according to their residue type (white, apolar; green, polar; blue, positively charged; red, negatively charged). In all panels, the homology model of mGBP7 is shown as a gray cartoon. In mGBP7$_{GTP}$, GTP and $Mg^{2+}$ are shown in green and orange.

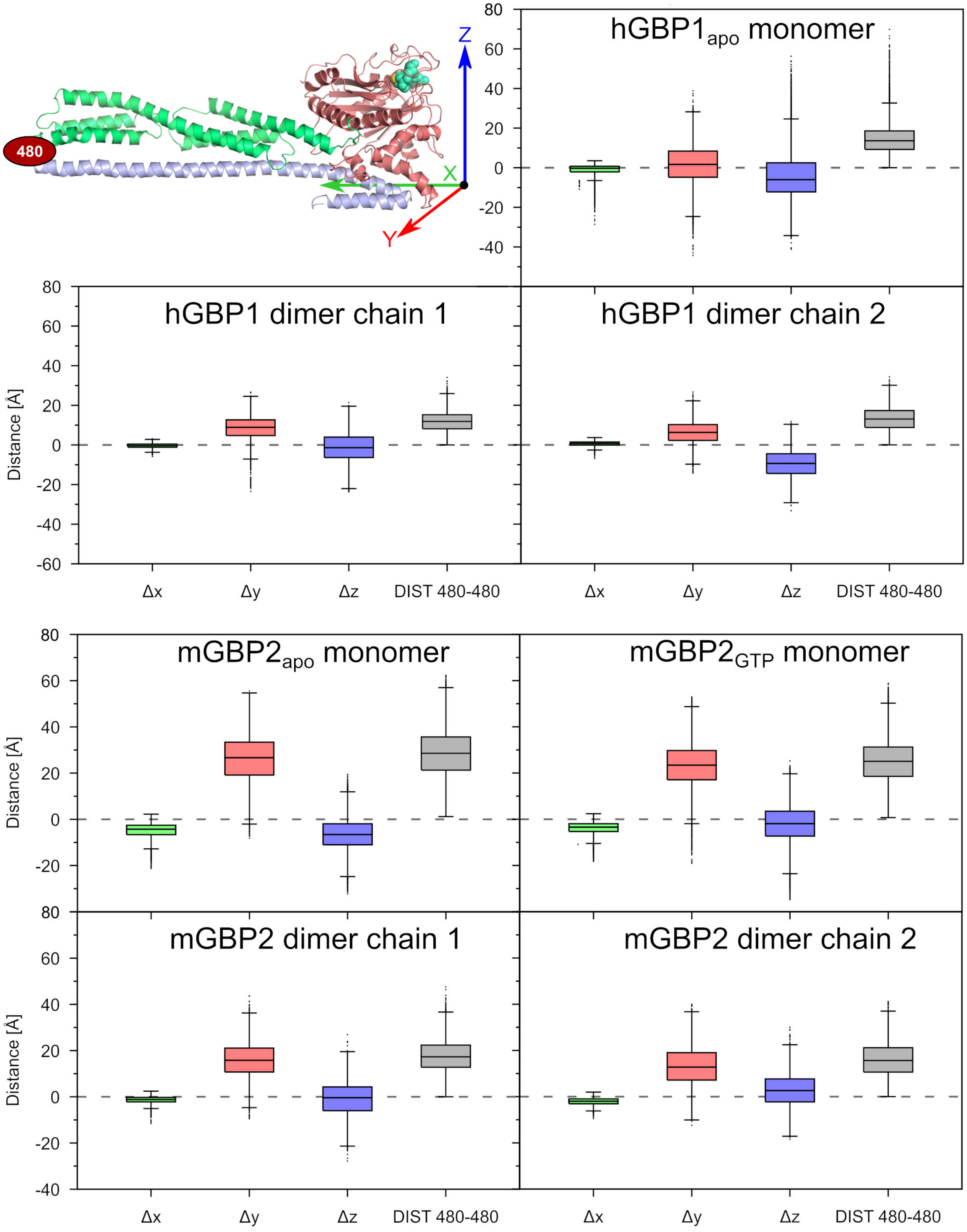


**Figure S5: Boxplots of $\Delta x$, $\Delta y$, and $\Delta z$ and $d_{480}$ of residue 480 for the hGBP1 and mGBP2 systems.** The orientation of the coordinate system used for these calculations is shown in the top left panel. The colors of the boxes correspond to the colors of the coordinate axes: $\Delta x$, green; $\Delta y$, red; $\Delta z$, blue. The $d_{480}$ boxes are shown in gray. The monomers and individual chains of the dimers were aligned to the hGBP1 or mGBP2 reference structure, illustratively shown for hGBP1 in the top left panel.

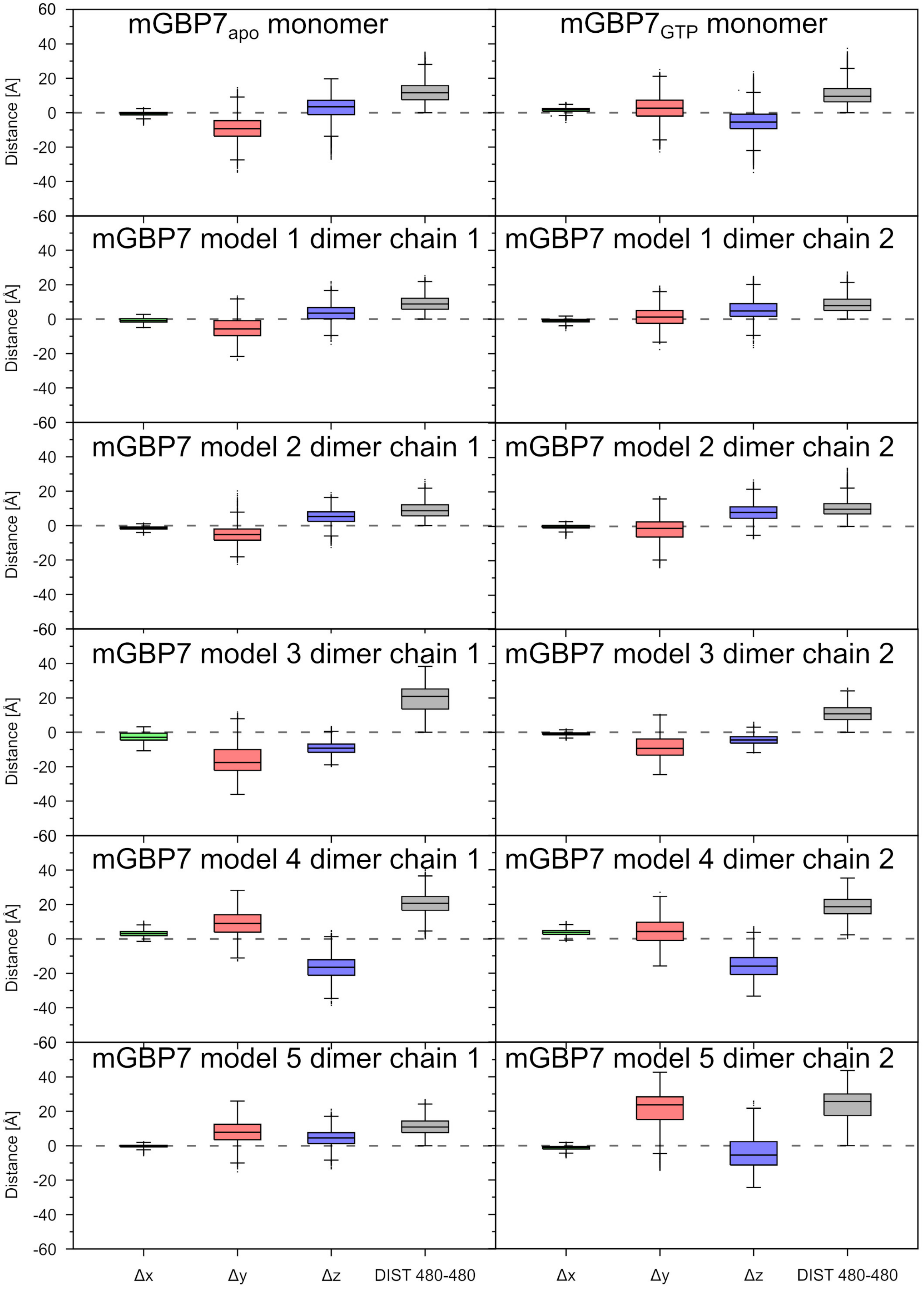


**Figure S6: Boxplots of $\Delta x$, $\Delta y$, and $\Delta z$ and $d_{480}$ of residue 480 for the mGBP7 systems.** See Fig. S5 for further details.

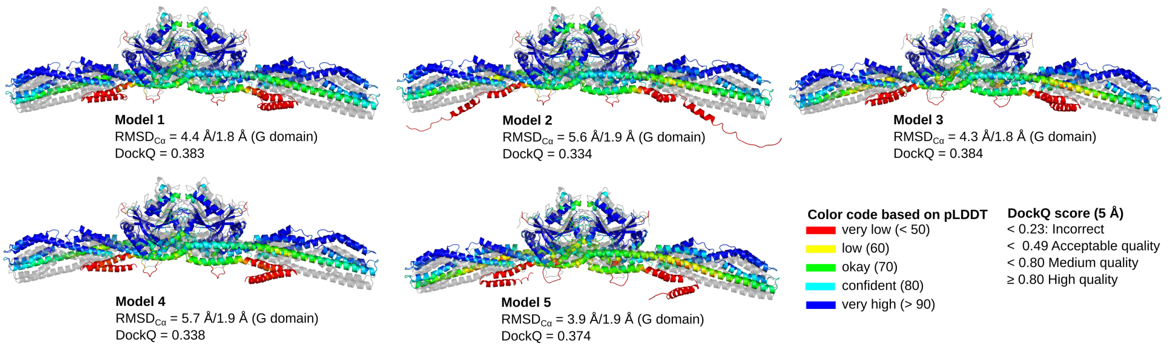


**Figure S7: The five most likely mGBP7 dimer models as predicted by AlphaFold-Multimer.** The dimer models are colored based on a per-residue estimate of the prediction's confidence (called pLDDT) on a scale from 0–100. Regions with pLDDT $> 90$ are shown in blue and expected to be modeled to high accuracy. Regions with pLDDT between 70 and 90 are expected to be modeled well (a generally good backbone prediction), while the predictions for regions with pLDDT between 50 and 70 are of low confidence and should be treated with caution. The 3D coordinates of regions with pLDDT $< 50$ should not be interpreted. Such low pLDDT values are a strong predictor of disorder or that the region in question is only structured as part of a complex. Most parts of the G and M domains in the AlphaFold-Multimer models are predicted with high confidence, while it is lower for $\alpha 12/13$ of the E domain and very low for the CT tail. The deviation from the G domain dimer model of mGBP2 (shown as gray cartoon) is provided in terms of the RMSD of the $C_\alpha$ atoms and the DockQ score.[27]

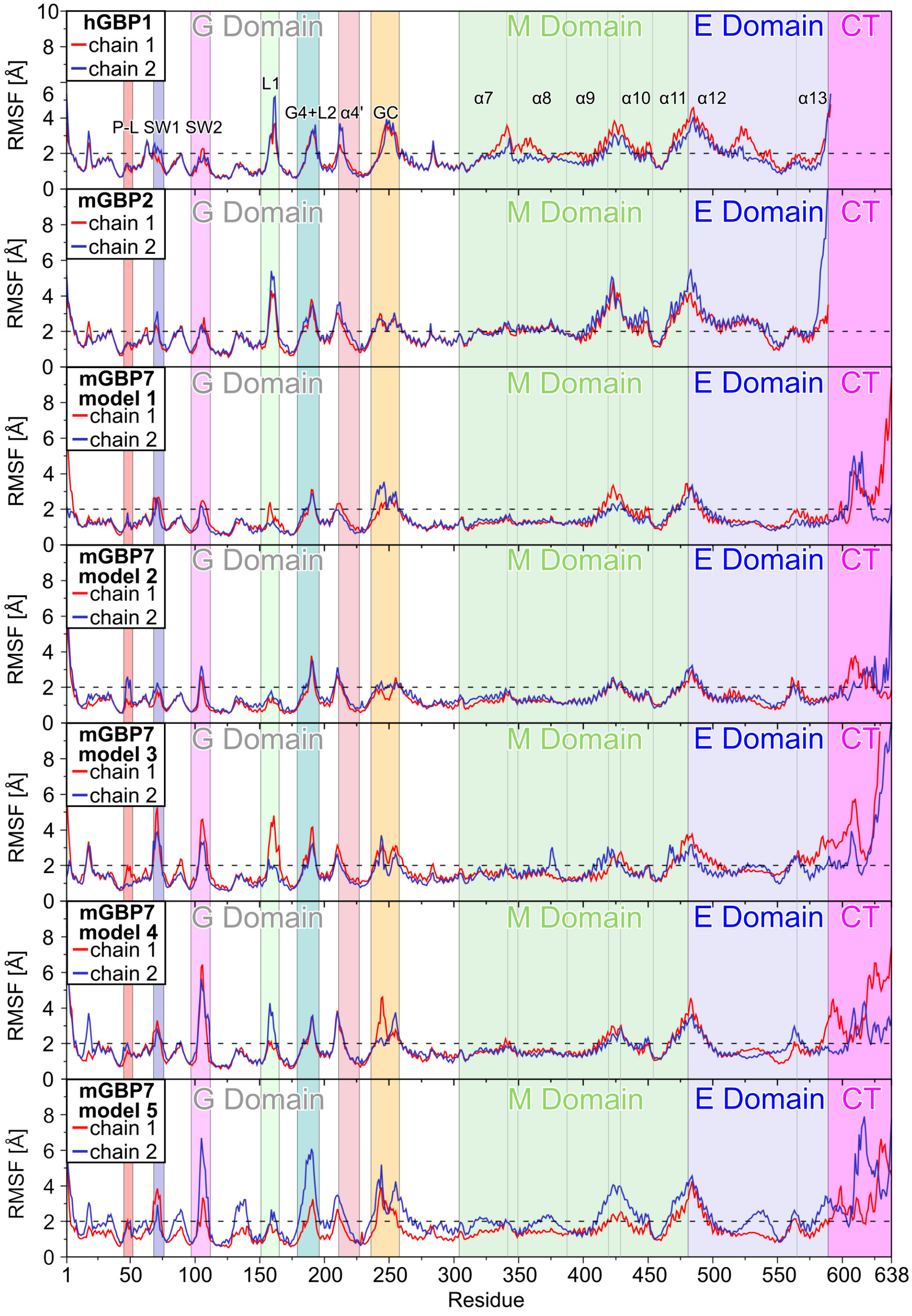


**Figure S8: Fluctuations of the GBP residues during MD simulations of the protein dimers.** The fluctuations are quantified by the RMSF of the $C_\alpha$ atoms of the two chains per dimer (shown in red and blue) after individually aligning the chains. All motifs and loops of the G domains as well as helices in the M/E domain are labeled, and the background of the plots is colored to indicate the different structural parts of the GBPs (where the same colors as in Fig. 1C/D were used). The horizontal dashed line at 2 Å is to identify flexible residues with RMSF values exceeding that value.

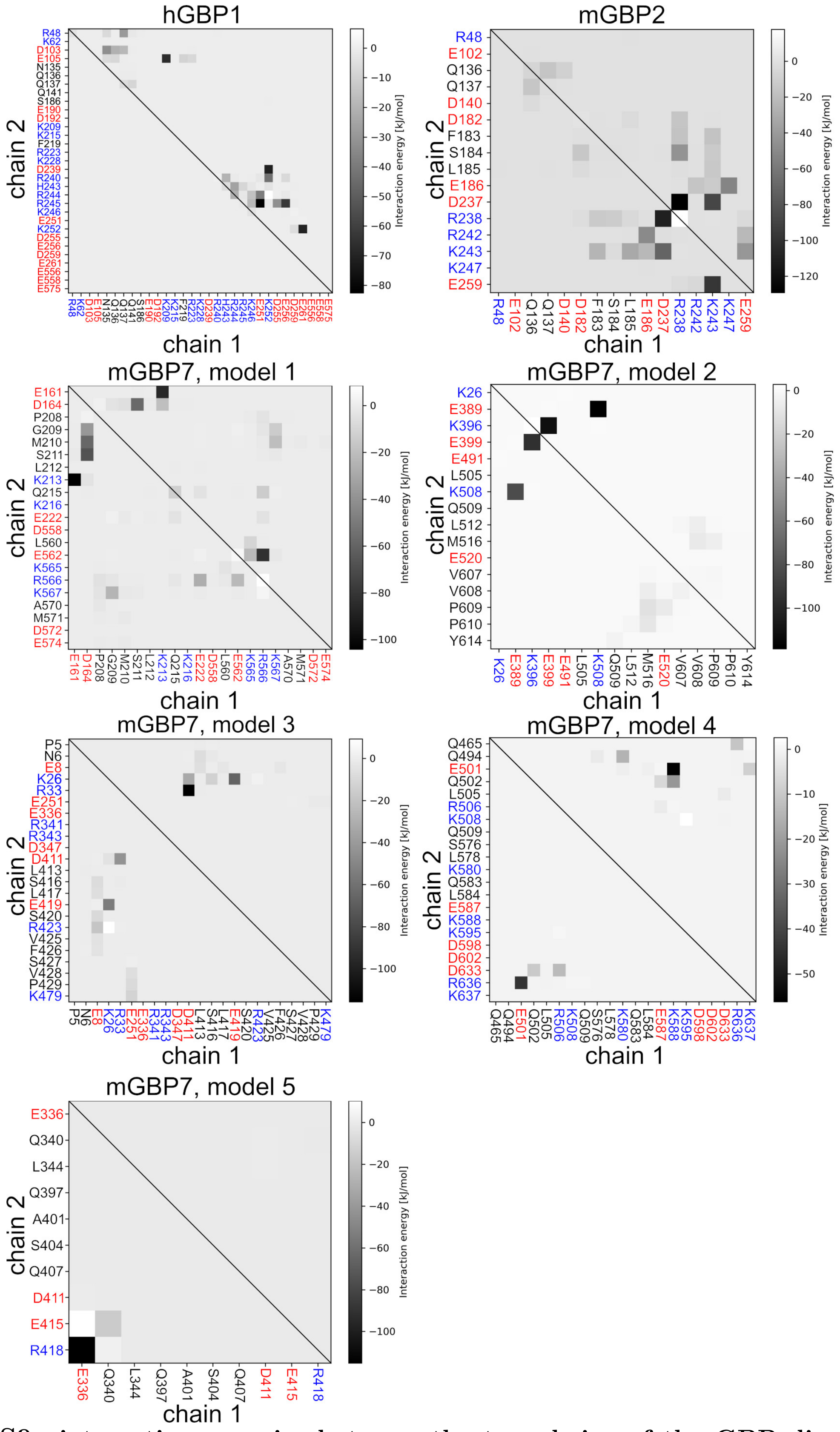


**Figure S9: interaction energies between the two chains of the GBP dimers.** These energies were calculated from the 100 ns MD simulations of the dimer models. Only the energies considerably different from zero are shown. The energies are given in kJ/mol, according to the color scale on the right per plot. It should be noted that the energie scales differ between the plots, as the color black was chosen for the minimal energy encountered per system. The residue labels are given on the axes, with red and blue for negatively and positively charged residues, respectively.

### 3 Supplementary information tables

**Table S1: Flexibility of the motifs and loops of the G domain and helix $\alpha$4'.** This analysis based on conformational clustering applied to the $C_\alpha$ atoms after aligning the target replica of the respective HREMD simulation on the $\beta$-sheets of G domain and using a 2.5 Å RMSD cutoff for asigning cluster membership.

| | **hGBP1$_{apo}$** | | |
|---|---|---|---|
| | **clusters** | **population [%]**[a] | **RMSD [Å]**[b] |
| P-L | 6 | 99.7 | 6.3/5.9 |
| SW1 | 93 | 61.7 | 14.8/13.4 |
| SW2 | 59 | 73.7 | 13.1/11.3 |
| L1 | 36 | 79.0 | 9.2/8.2 |
| G4+L2 | 55 | 64.4 | 12.4/11.8 |
| $\alpha$4' | 2 | 100 | 4.6/2.5 |
| GC | 197 | 34.4 | 20.4/19.9 |

| | **mGBP2$_{apo}$** | | | **mGBP2$_{GTP}$** | | | **%change**[c] |
|---|---|---|---|---|---|---|---|
| | **clusters** | **population [%]**[a] | **RMSD [Å]**[b] | **clusters** | **population [%]**[a] | **RMSD [Å]**[b] | |
| P-L | 1 | 100 | 4.4/– | 1 | 100 | 1.9/– | 0 |
| SW1 | 23 | 82.4 | 9.9/9.9 | 1 | 100 | 3.0/– | -95.7 |
| SW2 | 3 | 100 | 5.3/4.9 | 1 | 100 | 4.4/– | -66.7 |
| L1 | 23 | 89.6 | 11.2/10.8 | 17 | 92.2 | 9.9/9.4 | -26.1 |
| G4+L2 | 12 | 95.9 | 9.4/7.7 | 3 | 100 | 5.4/4.5 | -75.0 |
| $\alpha$4' | 2 | 100 | 5.4/3.4 | 3 | 100 | 6.1/5.2 | +33.3 |
| GC | 29 | 66.1 | 10.7/8.9 | 3 | 100 | 6.6/5.3 | -89.7 |

| | **mGBP7$_{apo}$** | | | **mGBP7$_{GTP}$** | | | **%change**[c] |
|---|---|---|---|---|---|---|---|
| | **clusters** | **population [%]**[a] | **RMSD [Å]**[b] | **clusters** | **population [%]**[a] | **RMSD [Å]**[b] | |
| P-L | 21 | 82.4 | 20.5/20.2 | 1 | 100 | 4.3/– | -95.2 |
| SW1 | 45 | 68.4 | 29.8/27.5 | 14 | 89.7 | 9.8/8.5 | -68.9 |
| SW2 | 68 | 44.0 | 35.1/34.6 | 32 | 78.1 | 12.6/10.9 | -52.9 |
| L1 | 100 | 54.4 | 35.6/34.5 | 30 | 77.3 | 11.4/9.6 | -70.0 |
| G4+L2 | 36 | 67.8 | 24.9/23.7 | 6 | 99.6 | 8.3/6.4 | -83.3 |
| $\alpha$4' | 63 | 58.2 | 36.0/34.4 | 17 | 92.3 | 8.8/8.5 | -73.1 |
| GC | 111 | 51.7 | 23.7/22.9 | 96 | 58.5 | 18.2/17.1 | -13.5 |

[a] Percentage of the structures which are cumulatively represented by the first three clusters.
[b] The largest RMSD found between any two clusters.
[c] $\%\text{change} = -(1 - \#\text{clusters(with GTP)}/\#\text{clusters(apo)}) \cdot 100$ or
$\%\text{change} = +(1 - \#\text{clusters(apo)}/\#\text{clusters(with GTP)}) \cdot 100$

**Table S2:** Summary of crosslinks by DSSO and BS3 for the mGBP7 monomer and dimer.

| State[a] | Residue pair[b] | Total[c] | model 1[d] | model 2[d] | model 3[d] | model 4[d] | model 5[d] |
|---|---|---|---|---|---|---|---|
| | | | **DSSO** | | | | |
| M & D | 106–216 | 5 | both | intra | intra | intra | intra |
| M & D | 495–440* | 5 | intra | intra | intra | intra | intra |
| M & D | 580–588 | 12 | intra | intra | intra | intra | intra |
| M & D | 586–539 | 7 | intra | intra | intra | intra | intra |
| M & D | 588–539* | 5 | intra | intra | intra | intra | intra |
| M & D | 595–524* | 8 | intra | both | intra | both | intra |
| (M) & D | 588–155 | 3 | intra | intra | intra | intra | intra |
| (M) & D | 205–554 | 4 | both | intra | intra | intra | intra |
| (M) & D | 205–565 | 4 | both | intra | intra | intra | intra |
| D | 373–588 | 2 | intra | intra | intra | intra | intra |
| D | 613–510 | 2 | intra | both | intra | both | intra |
| D | 613–524 | 2 | intra | both | intra | both | intra |
| D | 524–588* | 2 | intra | intra | intra | both | intra |
| D | 557–565 | 6 | both | intra | intra | intra | intra |
| | | | **BS3** | | | | |
| M & D | 588–539* | 6 | intra | intra | intra | intra | intra |
| M & D | 92–89 | 18 | intra | intra | intra | intra | intra |
| M & D | 216–216 | 18 | inter | no | no | no | no |
| M & D | 495–440* | 7 | intra | intra | intra | intra | intra |
| M & D | 611–510 | 5 | intra | both | intra | both | intra |
| M & D | 510–500 | 19 | intra | both | intra | intra | intra |
| (M) & D | 524–588* | 4 | intra | both | intra | both | intra |
| (M) & D | 567–565 | 4 | both | intra | intra | intra | intra |
| D | 565–565 | 2 | inter | no | no | no | no |
| D | 595–524* | 3 | intra | intra | intra | both | intra |
| D | 205–567 | 6 | both | intra | intra | intra | intra |
| D | 205–576 | 2 | both | intra | intra | intra | intra |
| D | 567–567 | 10 | inter | no | no | no | no |
| D | 567–576 | 2 | both | intra | intra | intra | intra |
| M | 98–89 | 2 | — | — | — | — | — |

[a] The crosslinks have been detected in the monomer (M) and/or dimer (D) band from crosslinked and size-separated mGBP7. Brackets are used if the linked peptide pair was found only in one of two experiments.
[b] Linked residues pairs found with both crosslinkers are marked by *.
[c] The total number of crosslinks for this residue pair in the monomer and/or dimer.
[d] It is indicated which crosslinks agree with the respective dimerization model of mGBP7: within a monomer (intra), intermolecular (inter), both intra- and intramolecular (both), no crosslink possible (no).

#### Table S3: Overall SAXS Data.

| SAXS Device | Xenocs Xeuss 2.0 with Q-Xoom |
|---|---|
| **Data collection parameters** | |
| Detector | PILATUS 3 R 300K windowless |
| Detector distance (m) | 0.550 |
| Beam size | 0.8 mm x 0.8 mm |
| Wavelength (nm) | 0.154 |
| Sample environment | Low Noise Flow Cell, 1 mm ø |
| s range ($nm^{-1}$)‡ | 0.05 – 6.5 |
| Exposure time per frame (s) | 600 (6 frames) |
| **Sample** | **mGBP7** |
| Organism | Mus musculus (Mouse) |
| UniProt ID | Q91Z40 (1-638) |
| Mode of measurement | batch |
| Temperature (°C) | 10 |
| Protein concentration (mg/ml) | 1.66 – 6.85 (merged) |
| Buffer | 50mM Tris pH 8.0, 5 mM MgCl, 2 mM DTT |
| **Structural parameters** | |
| I(0) from P(r) | 0.01 |
| $R_g$ (real-space from P(r)) (nm) | 5.08 |
| I(0) from Guinier fit | 0.01 |
| s-range for Guinier fit ($nm^{-1}$) | 0.099 – 0.240 |
| $R_g$ (from Guinier fit) (nm) | 5.35 |
| points from Guinier fit | 2 – 26 |
| $D_{max}$ (nm) | 17.00 |
| POROD volume estimate ($nm^3$) | 176.37 |
| **Molecular mass (kDa)** | |
| From I(0) | 137.09 |
| From Qp[28] | 136.31 |
| From MoW2[29] | 129.49 |
| Bayesian Inference[30] | 130.86 |
| From POROD | 110.23 |
| From sequence | 73.83 (monomer) |
| | 147.66 (dimer) |
| **Structure Evaluation** | |
| GASBOR fit $\chi_2$ | 1.14 |
| SASREF fit $\chi_2$ | 1.27 |
| Ambimeter score | 2.601 |
| Crysol fit $\chi_2$ | 1.89 |
| **Software** | |
| ATSAS Software Version[16] | 3.0.3 |
| Primary data reduction | PRIMUS[17] |
| Data processing | GNOM[19] |
| *Ab initio* modeling | GASBOR[20] |
| Rigid body modeling | SASREF[21] |
| Superimposing | SUPCOMB[22] |
| Structure evaluation | AMBIMETER[31] / CRYSOL[23] |
| Model visualization | PyMOL[32] |

‡ s = $4\pi sin(\theta)/\lambda$, $2\theta$ – scattering angle, $\lambda$ – X-ray-wavelength

**Table S4:** $R_g$ and $D_{max}$ of the mGBP7 dimer models in comparison to the SAXS data determined with CRYSOL.

| Protein | $\chi^2$ | $R_{g,sim}$ [nm] | $R_{g,SAXS}$ [nm] | $D_{max,sim}$ [nm] | Envelope diameter [nm] |
|---|---|---|---|---|---|
| model 1 | 1.66 | 5.5 | 5.50 | 23.1 | 24.1 |
| model 2 | 1.79 | 5.0 | 4.92 | 17.6 | 17.9 |
| model 3 | 1.74 | 4.9 | 4.97 | 14.9 | 16.7 |
| model 4 | 1.89 | 4.9 | 5.03 | 15.9 | 17.3 |
| model 5 | 1.27 | 5.5 | 5.67 | 18.6 | 19.1 |

(32) PyMol, The PyMOL Molecular Graphics System, Version 2.0. 2015; Schrödinger, LLC.
